# Nanoscale phosphoinositide distribution on cell membranes of mouse cerebellar neurons

**DOI:** 10.1101/2022.07.22.501145

**Authors:** Kohgaku Eguchi, Ryuichi Shigemoto

## Abstract

Phosphatidylinositol-4,5-bisphosphate (PI(4,5)P_2_) plays an essential role in neuronal activities through interaction with various proteins involved in signaling at membranes. However, the distribution pattern of PI(4,5)P_2_ and the co-clustering with these proteins on the neuronal cell membranes remain elusive at the electron microscopic level. In this study, we quantitatively investigated the nanoscale distribution of PI(4,5)P_2_ on the neuronal cell membranes by SDS-digested freeze-fracture replica labeling with cryo-fixed mouse cerebellum. We demonstrate that PI(4,5)P_2_ makes clusters with a mean size of ∼ 1,000 nm^2^ and these clusters show preferential accumulation in specific membrane compartments depending on cell types. Purkinje cell (PC) spines and granule cell (GC) presynaptic active zones are particularly rich in the PI(4,5)P_2_ clusters. Furthermore, these clusters are extensively associated with clusters of Ca_V_2.1 and GIRK3 throughout different membrane compartments of PCs, GCs, and molecular layer interneurons. In contrast, these clusters showed compartment-specific association with mGluR1α in PC spines. These results suggest that visualization of the nanoscale PI(4,5)P_2_ distribution may provide insight into the physiological functions of PI(4,5)P_2_ in neurons.

## Introduction

Phosphoinositides (PIs) are minor components on the cytoplasmic side of eukaryotic cell membranes, but they play essential roles in a wide variety of cellular functions. In neuronal cells, each stereoisomer of PIs is distributed in different subcellular compartments (Haucke, 2005; Idevall-Hagren and De Camilli, 2015; Wenk and De Camilli, 2004): PI(4)P is enriched in the membrane of the Golgi apparatus and synaptic vesicles (SVs), PI(4,5)P_2_ and PI(3,4,5)P_3_ mainly exist in the plasma membrane, PI(3)P and PI(3,5)P_2_ are selectively concentrated on early and late endosomes, respectively. PIs contribute to various aspects of neuronal activity, such as synaptic transmission and maintenance of membrane excitability by regulating ion channels and intracellular signaling pathways. At chemical synapses, PIs regulate exo- and endocytosis of synaptic vesicles at the presynaptic sites (Lei et al., 2017; Martin, 2001; Posor et al., 2015; Ueda and Hayashi, 2013)Many proteins related to SV exo- and endocytosis have binding domains to one or several stereoisomers of PIs (Di Paolo and De Camilli, 2006; Falkenburger et al., 2010), indicating a potential role of PIs as an anchor for these presynaptic proteins to regulate their localization and dynamics during synaptic transmission. At the postsynaptic sites, PIs regulate the morphological plasticity of dendritic spines through actin remodeling (Lei et al., 2017; Martin, 2001; Posor et al., 2015; Ueda and Hayashi, 2013)during long-term potentiation (Lei et al., 2017; Martin, 2001; Posor et al., 2015; Ueda and Hayashi, 2013).

Although it is essential to know the distribution pattern of PIs and their association with signaling proteins at neuronal cell membranes to understand the roles of PIs in neuronal activities, it has been poorly investigated due to several technical issues in the conventional methods (Tsuji et al., 2019). Immunoelectron microscopy using anti-PI(4,5)P_2_ antibody allows for observing the nanoscale PI(4,5)P_2_ distribution on the cell membrane, but the use of aldehyde-fixed cell preparations has a potential issue because aldehydes cannot fix the lateral diffusion of most membrane lipids across the membrane (Tanaka et al., 2010). Fluorescent probes for PIs based on a PI-binding domain (PBD) of proteins have been developed to visualize PIs in living cells (Idevall-Hagren and De Camilli, 2015; Maekawa and Fairn, 2014). However, this method does not have sufficient spatial resolution to observe the nanoscale distribution of PIs in small compartments in neuronal cell membranes, such as presynaptic active zones (AZs) and postsynaptic densities (PSDs). In addition, the overexpressed PBD-based probes mask PIs and competitively interfere with their interactions with proteins in the living cells (Suh and Hille, 2008).

To solve these issues and visualize the nanoscale distribution of PIs on cell membranes, sodium dodecyl sulfate-digested freeze-fracture replica labeling (SDS-FRL) combined with cryofixation has been utilized (Aktar et al., 2017; Cheng et al., 2014; Fujita et al., 2009; Ozato-Sakurai et al., 2011; Tsuji et al., 2019). This immunoelectron microscopic method enables nanoscale membrane phospholipid labeling by physically fixing membrane molecules without chemical fixation. In this study, we optimized this SDS-FRL method to acute mouse cerebellar slices and investigated the nanoscale distribution of PI(4,5)P_2_ on neuronal cell membranes using recombinant GST-tagged pleckstrin homology (PH) domain of phospholipase Cδ1 (PLCδ1) as a specific probe for PI(4,5)P_2_. This approach allowed us to quantitatively examine the numbers, densities, and distribution patterns of PI(4,5)P_2_ on somatodendritic and axonal membranes, including post- and presynaptic sites in mouse cerebellar neurons. We show that PI(4,5)P_2_ makes small clusters on neuronal membranes and specifically co-clusters with P/Q-type voltage-gated calcium channels, G-protein-coupled inwardly rectifying potassium channels, and metabotropic glutamate receptors in distinct membrane compartments, giving insights into the physiological functions of PI(4,5)P_2_ in the regulation of neuronal excitability and neurotransmitter release.

## Materials and Methods

### Animals

Animal experiments were conducted in accordance with the guideline of the Institute of Science and Technology Austria (Animal license number: BMWFW-66.018/0012-WF/V/3b/2016). Mice were initially purchased from Jackson Laboratory (Bar Harbor, ME, USA) and were bred at the Preclinical Facility of IST Austria on 12:12 light-dark cycle with access to food and water *ad libitum*. All experiments were performed in the light phase of the cycle. C57BL/6J (stock number 000664) mice of either sex at postnatal (P) 5-7 weeks were used in this study.

### Liposome preparation and high-pressure freezing

Phosphatidylcholine (18:1 (Δ9-Cis) PC), phosphatidylethanolamine (18:1 (Δ9-Cis) PE), phosphatidylserine (18:1 (Δ9-Cis) PS), phosphatidylinositol (18:1 PI) and phosphoinositides (18:1) were from Avanti Polar Lipids. All liposomes contained 45 mol % PC, 30 mol % PE, 20 mol % PS, and either 5 mol % PI or a phosphoinositide. Solutions of PC, PE, and PS in chloroform and PI or phosphoinositides in chloroform:methanol:H2O:HCl (1N) (20:9:1:0.1) were mixed in the required proportion in amber-color glass vials. To improve phosphoinositide homogenization with other lipids, the chloroform:methanol 2:1 ratio was maintained in the mixture. A lipid film was produced by evaporation of solvents in the vial under the stream of nitrogen gas and then drying using a vacuum desiccator for 2 h. The dried lipid film was stored at -20 °C with argon gas and used within 2 days.

The lipid film was resuspended in a buffer containing 220 mM sucrose and 20 mM HEPES (pH 7.4, adjusted with NaOH). The suspension was vortexed well and then freeze-thawed 5 times in liquid nitrogen and warm water (∼60 °C). Unilamellar liposomes were produced by extrusion through a 0.4 µm pore size polycarbonate filter using an extrusion apparatus (Avanti Polar Lipids). After extrusion, the liposomes were diluted 5 times in 120 mM NaCl, 20 mM HEPES buffer (pH 7.4, adjusted with NaOH), and centrifuged for 20 min at 15,000 g. The same amount of 100% glycerol was applied to the pellets of the liposomes as a cryoprotectant. The liposome/glycerol mixture was placed on a copper carrier with a ring of double-sided tape (140-µm thickness), covered with another carrier, and then frozen by a high-pressure freezing machine (HPM010, BAL-TEC). The frozen samples were stored in liquid nitrogen until use.

### High-pressure freezing of acute cerebellar slices

Acute slices of mouse cerebellum were prepared at physiological temperature (PT) as described previously (Eguchi et al., 2020). Briefly, mice were decapitated under isoflurane anesthesia and their brains were quickly removed from the skull and immersed into a cutting solution containing (mM): 300 sucrose, 2.5 KCl, 10 glucose, 1.25 NaH_2_PO_4_, 2 Na Pyruvate, 3 *myo*-inositol, 0.5 Na ascorbate, 26 NaHCO_3_, 0.1 CaCl_2_, 6 MgCl_2_ (pH 7.4 when gassed with 95% O_2_/5% CO_2_) at PT (35-37 °C). The cerebellum was dissected from the whole brain and immediately glued on a cutting stage of a tissue slicer (Linear Slicer Pro7, Dosaka EM, Kyoto, Japan) and sliced (sagittal, 140-160 µm thickness) in the cutting solution kept at PT. Slices were then maintained in the artificial cerebrospinal fluid (ACSF) containing (in mM): 125 NaCl, 2.5 KCl, 10 glucose, 1.25 NaH_2_PO_4_, 2 sodium pyruvate, 3 *myo*-inositol, 0.5 sodium ascorbate, 26 NaHCO_3_, 2 CaCl_2_, 1 MgCl_2_ (pH 7.4 when gassed with 95% O_2_/5% CO_2_) at 37 °C until use. Small blocks containing lobule IV-VII were trimmed from the slices in the cutting solution using a micro scalpel (#10316-14, FST). The trimmed block was transferred into cryoprotectant buffer (15% polyvinylpyrrolidone in ACSF with 10 mM HEPES, pH 7.3 adjusted with NaOH), sandwiched between two copper carriers with a ring of double-sided tape (140-µm thickness), and then frozen by a high pressure freezing machine. The frozen samples were stored in liquid nitrogen until use.

### SDS-digested freeze-fracture replica labeling (SDS-FRL)

The frozen samples were fractured into two parts at -130 °C and replicated by carbon (4-5 nm thick), carbon-platinum (uni-direction from 60°, 2 nm), and carbon (20-25 nm) deposition in a freeze-fracture machine (JFD-V, JOEL, Tokyo, Japan). The samples were digested with 2.5% SDS solution containing 0.1 M Tris-HCl (pH 8.3) at 80°C for 18-22 h. The replicas were washed in the SDS solution and then a washing buffer (50 mM Tris-buffered saline (TBS, pH 7.4) containing 0.1% BSA) at RT. To avoid non-specific binding of the probes and antibodies, the replicas were blocked with 3% BSA, 2% cold fish skin gelatin, and 0.05% Tween-20 in TBS for 1 h at RT. The replicas were incubated with 50 ng/ml GST-tagged PH domain of phospholipase C δ1 (PI(4,5)P_2_-Grip, Echelon Inc.) in a dilution buffer (1% BSA, 1% cold fish skin gelatin, and 0.05% Tween-20 in TBS) at 4°C overnight. Then the replica was incubated with anti-GST antibody (rabbit IgG, 5 µg/ml, Bethyl Laboratories) and anti-Ca_V_2.1 antibody (guinea pig, 2.5 µg/ml, Synaptic Systems) as a marker of neurons at 15°C overnight, and then gold-nanoparticle conjugated secondary antibodies (goat anti-rabbit IgG, 5 nm, 1:50, BBI; donkey anti-guinea pig IgG, 12 nm, 1:30, Jackson Immnoresearch) dissolved in the dilution buffer at 15°C overnight. For double labeling of PI(4,5)P_2_ with proteins, the replicas were incubated with a mixture of primary antibodies (GluD2: rabbit anti-GST IgG + guinea pig anti-GluD2 IgG (2.0 µg/ml, Frontier Institute); GIRK3: chicken anti-GST IgY (5 µg/ml, Bethyl Laboratories) + rabbit anti-GIRK3 IgG (XX µg/ml, Frontier Institute); mGluR1α: rabbit anti-GST IgG + guinea pig anti-mGluR1α IgG (XX µg/ml, Frontier Institute)) at 15°C 1-2 overnight and then with gold-nanoparticle conjugated secondary antibodies at 15°C overnight. After washing the replicas with the washing buffer, they were picked up onto a grid coated with formvar in distilled water. Images were obtained under TEM (Tecnai 10) with RADIUS software at magnifications of 65,000 and 39,000.

### Image analysis

Images were analyzed with Darea software (Kleindienst et al., 2020), Fiji, and R. The gold particle detection and the demarcation of the region of interest were performed on Darea software. AZs on the P-face of PF boutons were indicated with the aggregation of intramembrane particles on the replica at the electron microscopic level as described previously (Eguchi et al., 2022, 2020; Harris and Landis, 1986; Landis and Reese, 1974; Masugi-Tokita et al., 2007). Gold particles inside or < 30 nm away from the demarcation border of AZs (outer rim) were counted as the particles in the AZs (Kleindienst et al., 2020). Because the PSD area on the P-face of dendritic spines of PCs cannot be identified based on morphological features, the largest cluster of GluD2-labeling gold particles on the spines was identified as the PSD area (Eguchi et al., 2020; Luján et al., 2018a). The demarcated region of the images was imported to R via FIJI/ImageJ for the following point pattern analysis. Point pattern analysis of the gold particles described below was performed using spatstat package (version 2.3-0) of R (version 4.1.0). Nearest neighbor distances (NNDs) to both particles of the same size (e.g. from a 5 nm particle to the nearest 5 nm particle) and the other size (e.g. from a 5 nm particle to the nearest 10 nm particle) were computed to evaluate the distribution pattern of the particles. Center Periphery Index (CPI), indicating the location of the particles in the AZs or PSDs, was calculated as the square of the normalized distances from the center of the region of interest. When particles are randomly distributed in a circle, the mean CPI is near 0.5 (Kleindienst et al., 2020). To assess the randomness of the particle distribution, we performed two types of Monte-Carlo simulations, termed random and fitted simulations, following the methods described in the previous publications (Kleindienst et al., 2020; Luján et al., 2018b) using R. For the random simulation, particles were randomly placed on the demarcated area. The simulated particles were placed to keep the minimum distance of 10 nm from any other particles and then randomly shifted within a disc with a 30-nm diameter to reproduce the immunolabeling with a probe and antibodies (Tabata et al., 2019). In the fitted simulation, a constraint was added that the distance distribution between the simulated particles should not differ significantly from the distance distribution between the original particles. The particle distribution pattern was modeled and simulated as a Matern Cluster point process, and the goodness-of-fit between the real and simulated distribution was assessed by comparing both the all pairwise distances (APD) and NNDs of the particles using the two-sample *Kolmogorov-Smirnov* test (KS test), respectively, and considered them similar if the p-value was equal or above 0.1 for both. To avoid excessive statistical power due to the large sample size caused by a large number of particles, parametric bootstrapping was performed when the number of values to be compared exceeded 100. Specifically, we first randomly selected 100 distance values (NND or APD) from the simulated distribution and compared them to the empirical cumulative distribution function of the distance values of the real distribution using the KS test. This process was repeated 1,000 times, and the average of the p-values was used to assess the goodness-of-fit. Gold particle clusters were detected with a hierarchical clustering algorithm called Ward linkage (Figure S1) and a density-based clustering algorithm called DBSCAN (Figures 2 and 3). For Ward linkage, the threshold distance was set as 50 nm. For DBSCAN, we set the minimum number of particles consisting of a cluster as 3 and the maximum distance between particles as the sum of the median and 1.5 times the interquartile range (IQR) of the NNDs. The area of the convex polygon connecting the outermost particles forming the cluster was defined as the cluster area.

### Statistical analysis

To consider the hierarchical structure, correlation, and probability distribution, data were analyzed with either a linear mixed-effects model (LMM) or its generalized form (GLMM) using the lme4 package (version 1.1-27.1) of R (Aarts et al., 2014; Yu et al., 2021). Probability distributions for models were chosen by the goodness of fit to Poisson (for discrete variables e.g. the number of particles), normal (e.g. CPI), or gamma distribution (for continuous variables e.g. NNDs). Appropriate to the particular experiment and statistical model, fixation methods (perfusion- and cryo-fixation), the faces of cell membranes (P- and E-face), components of neurons (e.g. somata, spines, AZs on PF boutons), genotypes (wildtype and Ts65Dn), and potentially their interactions were used as fixed effects, while experiments, animals, replicas, and cells were used as random effects to consider the nested and/or crossed data structure in the statistical analysis. For all experiments, at least four animals per condition were used. All data are presented as estimated marginal means with 95% confidence intervals (CI, in figures) or standard error of means (s.e.m., in text) estimated using the emmeans package. The goodness-of-fit of the models was assessed by second-order Akaike Information Criterion (AICc) for the elimination of the random factors (MuMIn package and then by likelihood ratio chi-square tests (Chi-LRT) with models in which the fixed effects of interest had been dropped. For multiple pairwise comparisons between 3 or more groups, posthoc comparisons to assume the significance of differences between pairs of group means were performed using emmeans packages with Tukey (for all pairwise) or Benjamini-Hochberg (BH, for selected pairwise) method when Chi-LRT detected a significant difference (p < 0.05). Statistical significance was assumed if p < 0.05 (indicated with blue or single asterisk), p < 0.01 (green or double asterisk), and p < 0.001 (red or triple asterisk).

## Results

### Visualization of nanoscale PI(4,5)P_2_ distribution on SDS-digested freeze-fracture replica preparations of mouse brain tissues

To observe the nanoscale two-dimensional distribution of PI(4,5)P_2_, SDS-FRL has been utilized on cultured human fibroblasts, mouse smooth muscle cells, and rat pancreatic exocrine acinar cells (Fujita et al., 2009; Ozato-Sakurai et al., 2011). In this study, we optimized this method for mouse cerebellar tissues to quantitatively analyze the nanoscale PI(4,5)P_2_ distribution on neuronal cell membranes. We used a recombinant GST-tagged PH domain of PLCδ1 (GST-PH) as a specific probe of PI(4,5)P_2_. To verify the specificity of GST-PH among the PI stereoisomers, we labeled freeze-fracture replicas of liposomes containing either PhdIns or a PI stereoisomer (5 mol%) with GST-PH (50 ng/ml), anti-GST primary antibody, and gold particle-conjugated secondary antibody. The density of immunogold particles was much higher on the replica prepared from PI(4,5)P_2_ liposome compared to those from other liposomes (Figure 1A), indicating a high specificity of the PI(4,5)P_2_ probe (Figure 1B).

**Figure 1.**
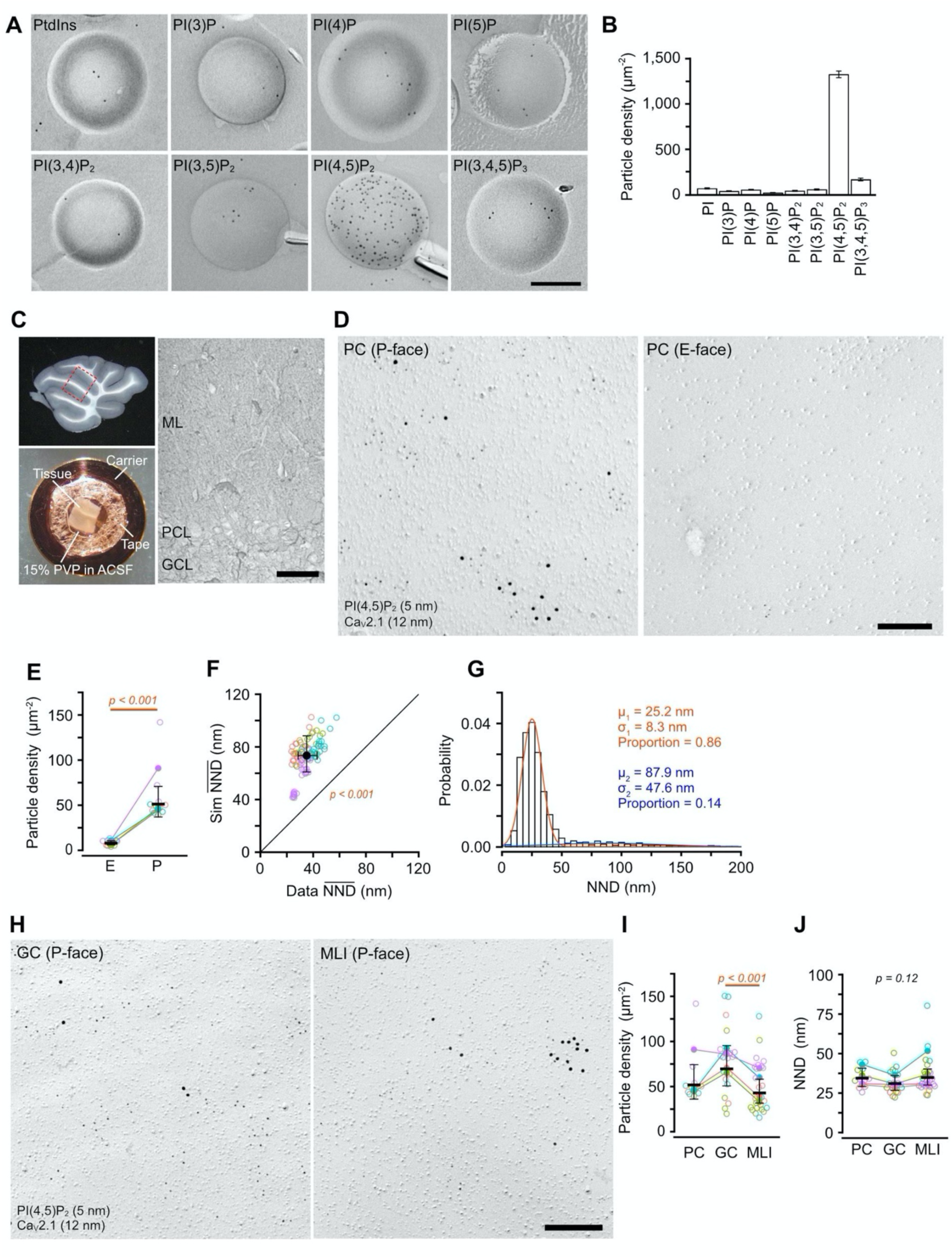
SDS-FRL of PI(4,5)P_2_ on somatic membranes of mouse cerebellar neurons. (**A-B**) Specificity of PI(4,5)P_2_ labeling. Replicas of liposomes containing 5% PtdIns or different stereoisomers of PIs were labeled using GST-PH, anti-GST antibody, and 5-nm gold particle-conjugated secondary antibody (**A**, Scale bar = 200 nm). The density of gold particles was highest in the liposome containing PI(4,5)P_2_ (**B**). (**C**) Acute cerebellar slice preparation for high-pressure freezing (HPF) and replica preparation. Left-top, an acute sagittal slice of the mouse cerebellum. The dashed line indicates the trimmed region for HPF. Left-bottom, a trimmed cerebellar slice on a copper carrier with double-sided tape for HPF. Right, low-magnification transmission electron microscopic (TEM) image of the mouse cerebellar replica containing granule cell layer (GCL), Purkinje cell layer (PCL), and molecular layer (ML). Scale bar = 20 µm. (**D**) Example TEM image of 5-nm gold particle labeling for PI(4,5)P_2_ with immunogold labeling for Ca_V_2.1 (12 nm) on P-(left) and E-face (right) of the PC somatic membranes of cerebellar PC. Scale bar = 200 nm. (**E**) Statistical comparison of the PI(4,5)P_2_ particle density on the E-face and P-face of the PC somatic membranes. Open and closed circles indicate the means of the PI(4,5)P_2_ particle density in each PC and each mouse, respectively, with different colors. Black horizontal bars indicate estimated marginal means (emmeans, thick bars) and 95% confidence intervals (CIs, error bars) of the density estimated by GLMM (Methods). The PI(4,5)P_2_ density was significantly higher on the P-face than on the E-face of the PC somatic membranes (P-face: 51.2 ± 8.5 particles/µm^2^, E-face: 8.0 ± 1.2 particles/µm^2^, n = 213 images/12 cells/4 mice, p < 0.001, Chi-square likelihood ratio test (Chi-LRT)). (**F**) Comparison of nearest neighbor distances (NND) between real (Data NND, x-axis) and simulated (Sim NND, y-axis) PI(4,5)P_2_ particles on PC somatic membranes. Data-NNDs are significantly smaller than Sim NNDs (Data: 35.1 ± 3.6 nm, Sim: 73.3 ± 7.0 nm, n = 206 images/11 cells/4 mice, p < 0.001, Chi-LRT). (**G**) Distribution of NNDs of the PI(4,5)P_2_ particles obtained from a single PC somatic membrane (n = 2,929 particles). Red and blue lines indicate the distinct components of the NND distribution estimated from the Gaussian mixture modeling. (**H**) PI(4,5)P_2_ labeling (5 nm) with Ca_V_2.1 (12 nm) on the P-face of GC (left) and molecular layer interneuron (MLI, right) somatic membranes. Scale bar = 200 nm. (**I**) The PI(4,5)P_2_ particle density on the somatic membranes (P-face) of different neuronal cell types in the mouse cerebellum (PC: 51.8 ± 9.6 particles/µm^2^, GC: 69.8 ± 11.3 particles/µm^2^, MLI: 42.9 ± 6.7 particles/µm^2^, n = 273 images/57 cells/4 mice, p = 0.003, Chi-LRT). (**J**) NND of the PI(4,5)P_2_ particles on the somatic membranes of different neuronal cell types in the mouse cerebellum (PC: 34.4 ± 3.0 nm, GC: 31.0 ± 2.4 nm, MLI: 34.8 ± 2.6 nm, n = 65,607 values/57 cells/4 mice, p = 0.12, Chi-LRT).

To visualize the two-dimensional (2D) distribution of PI(4,5)P_2_ on neuronal cell membranes, we labeled PI(4,5)P_2_ in the mouse cerebellum using SDS-FRL with GST-PH. Acute slices prepared at physiological temperature (Eguchi et al., 2020) were frozen under high pressure with 15% PVP as a cryoprotectant (Figure 1C) to minimize damage caused by ice crystals during freezing (Borges-Merjane et al., 2020). After the SDS-digestion to remove cytosolic proteins and extracellular matrix, cerebellar replicas were sequentially incubated with GST-PH, primary antibodies, and gold particle-conjugated secondary antibodies. Ca_V_2.1 was co-labeled using an anti-Ca_V_2.1 antibody directed against its intracellular domain (Althof et al., 2015), giving specific labeling on the cytoplasmic leaflets (P-face) of neuronal cell membranes. Immunogold particles that label PI(4,5)P_2_ by GST-PH (PI(4,5)P_2_ particles) showed higher density on the P-face of the somatic membrane of Purkinje cells (PCs) than on the E-face (Chi-square likelihood-ratio test (Chi-LRT): p < 0.001; Figures 1D and 1E), indicating the dominance of PI(4,5)P_2_ distribution on the cytoplasmic leaflet of the cell membranes. The particle density without GST-PH was significantly lower on both E- and P-face of the PC somatic membranes than that with GST-PH (Figures S1A and S1B), indicating that even the low-density PI(4,5)P_2_ labeling on the E-face is ascribable to GST-PH binding.

To quantitatively investigate whether PI(4,5)P_2_ is clustered on somatic membranes of PCs, we performed Monte-Caro random simulations and compared the nearest neighbor distance (NND) between the observed and simulated PI(4,5)P_2_ particles from all images (Szoboszlay et al., 2017). The mean NNDs of the observed PI(4,5)P_2_ particles were around half of the simulated one (Chi-LRT: p < 0.001, Figure 1F), indicating clustering of PI(4,5)P_2_ on the membrane. The distribution of NNDs between PI(4,5)P_2_ particles obtained from a PC somatic membrane (Figure 1G) is highly right-skewed (skewness = 3.55). Fitting this distribution with Gaussian mixture modeling shows that the NNDs between PI(4,5)P_2_ particles are produced from a mixture of short and long NND populations. These results indicate that PI(4,5)P_2_ on the PC soma has two distinct distribution patterns, clustered and scattered, and that most PI(4,5)P_2_ constitute clusters (Figure 1G). The PI(4,5)P_2_ particle clusters were also observed on the E-face of the somatic membrane (Figure S1C). The number of particles in a cluster was significantly lower without GST-PH than that with GST-PH on both E- and P-face (Figure S1C). These results indicate that PI(4,5)P_2_ forms clusters on both the outer and inner leaflets of the somatic membranes.

To examine whether the density of PI(4,5)P_2_ differs between neuronal cell types in the cerebellum, we observed somatic membranes of cerebellar granule cells (GCs) and molecular layer interneurons (MLIs). PI(4,5)P_2_ particles were also distributed on the P-face making clusters in somatic membranes of both cell types (Figure 1H). Although the density of PI(4,5)P_2_ particles in the somatic membrane of GCs was significantly higher than that of MLIs (multiple pairwise comparisons with Tukey adjustment (Tukey): p < 0.001; Figure 1I), the NNDs of the PI(4,5)P_2_ particles on somatic membranes of the cerebellar neurons did not differ between the cell types (Chi-LRT: p = 0.12; Figure 1J), indicating a similar local concentration of PI(4,5)P_2_ in these clusters across the examined cerebellar neuronal cell types.

### PI(4,5)P_2_ distribution on somatodendritic compartments of PCs

We next investigate whether the distribution of PI(4,5)P_2_ differs among somatodendritic compartments of PCs: somata, main shafts (MS), smooth branchlets (SmB), spiny branchlets (SpB), and spines. EM pictures of each dendritic component were taken in the molecular layer (ML) divided into proximal (MS), intermediate (SmB) and distal one-third (SpB, spines) of the PC layer. Gold particles for PI(4,5)P_2_ were observed throughout all the dendritic compartments (Figure 2A). The density of PI(4,5)P_2_ particles was found to increase gradually from the proximal to the distal dendrites (Chi-LRT: p < 0.001; Figure 2B): spine membranes showed a ∼1.5 times higher density of PI(4,5)P_2_ particles than somatic, MS, and SmB membranes (Tukey: p < 0.001), whereas no significant difference compared to SpB membranes was detected (Tukey: p = 0.43). Mean NNDs of PI(4,5)P_2_ particles were not significantly different between the compartments (Figure 2B, Chi-LRT: p = 0.55), indicating that the local concentration of PI(4,5)P_2_ does not differ between different somatodendritic compartments of PCs.

**Figure 2.**
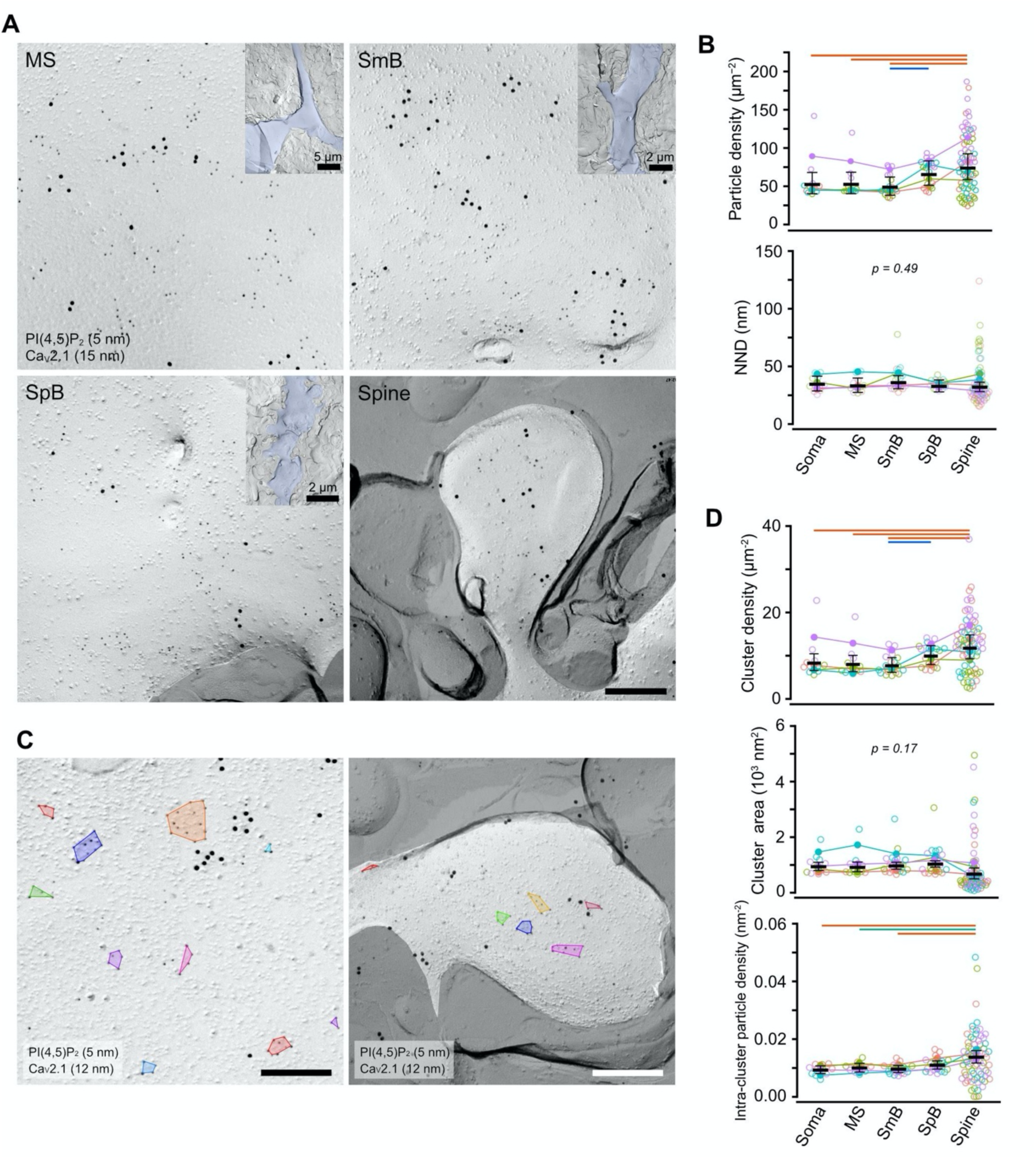
PI(4,5)P_2_ particle distribution on somatodendritic membranes of PCs. (**A**) Example images obtained from the different compartments of PC dendritic membranes: main shaft (MS, top-left), smooth branchlet (SmB, top-right), spiny branchlet (SpB, bottom-left), and spine (bottom-right). Scale bar = 200 nm. Inset: low-magnification images of MS, SmB, and SpB, respectively, indicated with blue. (**B**) Comparison of the PI(4,5)P_2_ particle density (top) and NNDs (bottom) in the somatodendritic membrane compartments of PCs. The PI(4,5)P_2_ density gradually increased from soma and proximal dendrites to distal dendritic components (top, n = 611 images/134 components/4 mice, p < 0.001, Chi-LRT), whereas no significant difference in NNDs was detected between these compartments (bottom, n = 67,380 values/134 components/4 mice, p = 0.49, Chi-LRT). (**C**) Example images of the PI(4,5)P_2_ clusters on the somatic (left) and spine (right) membrane of PCs. Colored polygons indicate individual PI(4,5)P_2_ clusters detected by the DBSCAN algorithm. Scale bars = 200 nm. (**D**) Quantitative analysis of the PI(4,5)P_2_ clusters on the somatodendritic compartments of PCs. Top, the number of clusters per area (cluster density, n = 611 values/134 components/4 mice). Middle, the cluster area (10,479 clusters/124 components/4 mice). Bottom, the intra-cluster particle density (10,343 clusters/134 components/4 mice). The cluster density and the intra-cluster particle density are significantly higher in spine membranes compared to the somatic and proximal dendritic membranes (p < 0.001, Chi-LRT), whereas there is no significant difference in the cluster area (p = 0.17, Chi-LRT).

PI(4,5)P_2_ particles on the dendritic membranes formed clusters as seen on the somatic membranes. To quantitatively compare the PI(4,5)P_2_ cluster profiles (cluster area, particle number in cluster, and intra-cluster density) between all compartments of somatodendritic membranes, we detected the clusters using the Density-Based Spatial Clustering of Applications with Noise (DBSCAN) algorithm (Methods, (Szoboszlay et al., 2017). The DBSCAN detected variable sizes of PI(4,5)P_2_ particle clusters on the somatodendritic membranes (Figure 2C). The density of PI(4,5)P_2_ clusters on the PC cell membranes gradually increased from the proximal to the distal compartments (Chi-LRT: p < 0.001; Figure 2D); spine membranes showed a higher density of PI(4,5)P_2_ clusters than somatic, MS, and SmB membranes (Tukey: p < 0.001), whereas no significant difference with SpB membranes was detected (Tukey: p = 0.21). The cluster area was not significantly different between all compartments of somatodendritic membranes (Chi-LRT: p = 0.10; Figure 2D), whereas intra-cluster PI(4,5)P_2_ particle density in spines was significantly higher than those in other compartments (Chi-LRT: p < 0.001; Figure 2D). These results suggest that PI(4,5)P_2_ are distributed throughout all compartments of somatodendritic membranes of PCs, and the density is higher at the distal dendrites and spines than at the soma and proximal dendrites. This proximo-distal gradient of PI(4,5)P_2_ along the somatodendritic compartments of PC membranes is ascribable to an increase in the density of PI(4,5)P_2_ clusters rather than the cluster size, and a higher PI(4,5)P_2_ density within clusters on spine membranes.

### PI(4,5)P_2_ distribution on membrane compartments of GCs and MLIs

We next examined the PI(4,5)P_2_ distributions in different compartments of GCs (somata, dendrites, parallel fiber (PF) axons, presynaptic PF boutons with PC spines (PF-PC) or MLI dendrites (PF-MLI)) and MLIs (somata, dendrites, presynaptic basket cell boutons on PC somata (BC-PC)). Gold particles for PI(4,5)P_2_ were observed throughout all compartments of GCs and MLIs (Figures 3A and 3B). In the GC membranes, the density of PI(4,5)P_2_ particles was not significantly different between the compartments (Chi-LRT: p = 0.095; Figure 3C), though the mean NNDs of PI(4,5)P_2_ particles were slightly but significantly different between somatic and axonal membranes (Tukey: p = 0.027, Figure 3C). The cluster parameters of PI(4,5)P_2_ were also not significantly different between the compartments (Figure 3D). These results suggest that PI(4,5)P_2_ in GC membranes shows a similar distribution pattern throughout the subcellular compartments.

**Figure 3.**
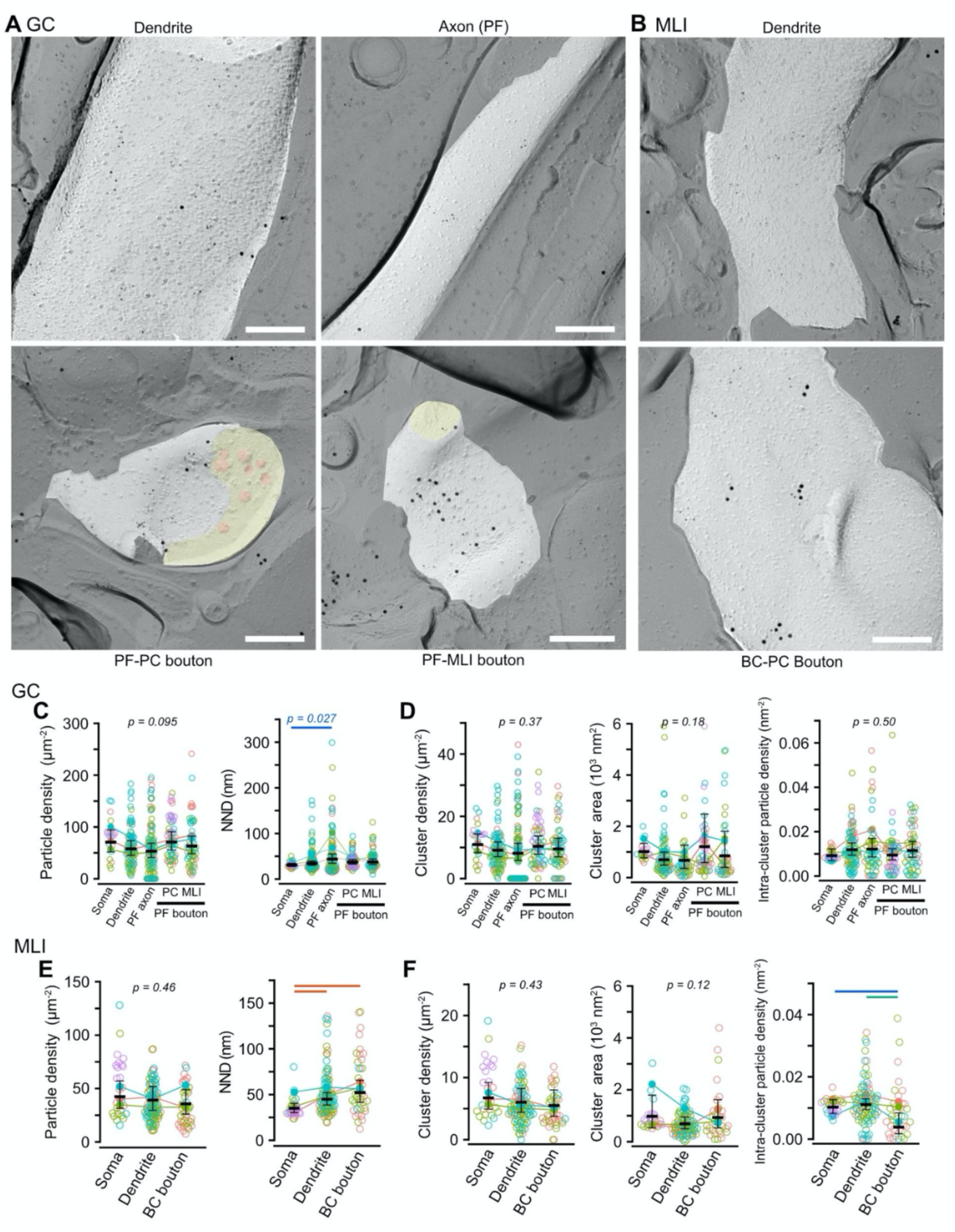
PI(4,5)P_2_ particle distribution in GCs and MLIs. (**A**) Example images obtained from the different compartments of GC membranes: dendrites (top-left), parallel fiber (PF) axons (top-right), presynaptic PF boutons on the PC spine (PC-PF, bottom-left) and on the MLI dendrite (PF-MLI, bottom-right). Yellow and orange in the PF boutons indicate the cross-fracture and synaptic vesicle membranes, respectively. Scale bars = 200 nm. (**B**) Example images obtained from different compartments of MLI membranes: dendrites (top) and basket cell (BC) presynaptic bouton on PC soma (bottom). Scale bar = 200 nm. (**C**) Comparison of the PI(4,5)P_2_ particle density (left) and NNDs (right) in the different GC membrane compartments. NNDs show significant differences between the compartments (n = 67,380 values/134 components/4 mice, p < 0.001, Chi-LRT), whereas there is no significance in the density (n = 376 images/320 components/4 mice, p = 0.09, Chi-LRT). (**D**) Quantitative analysis of the PI(4,5)P_2_ particle clusters in different GC membrane compartments. There are no significant differences in the cluster density (n = 376 images/320 components/4 mice, p = 0.37, Chi-LRT), the cluster area (n = 4070 clusters/254 components/4 mice, p = 0.18, Chi-LRT), and the intra-cluster particle density (n = 4004 clusters/253 compartments/4 mice, p = 0.50, Chi-LRT). (**E**) Comparison of the PI(4,5)P_2_ particle density (left) and NNDs (right) in different membrane compartments of MLIs. NNDs in dendritic and BC bouton membranes are significantly larger than that in somatic membranes (n = 24,012 values/152 components/4 mice, p < 0.001, Chi-LRT), whereas there is no significant difference in the density between the compartments (n = 230 images/153 components/4 mice, p = 0.46, Chi-LRT). (**F**) Quantitative analysis of the PI(4,5)P_2_ particle clusters in different MLI membrane compartments. The intra-cluster particle density in BC bouton membranes is significantly smaller than the other components (n = 3,766 clusters/149 components/4 mice, p = 0.03, Chi-LRT), whereas no significant difference was detected in the cluster density (n = 230 images/153 components/4 mice, p = 0.43, Chi-LRT) and the cluster area (n = 3,822 clusters/146 components/4 mice, p = 0.12, Chi-LRT) between these compartments.

In MLIs, the PI(4,5)P_2_ density was not significantly different between the membrane compartments, but the mean NNDs were larger in dendritic and presynaptic bouton membranes than in somatic membranes (Figure 3E). This difference is caused by larger population of scattered PI(4,5)P_2_ in these compartments as shown in Figure S2. The intra-cluster density of PI(4,5)P_2_ in the bouton membrane was significantly higher than in the somatic membrane, whereas the cluster density and area were similar between the compartments (Figure 3F).

### PI(4,5)P_2_ distribution on synaptic membranes

Next, we focused on PI(4,5)P_2_ distribution in the presynaptic membrane of PF (Figure 4A). Since many presynaptic proteins related to neurotransmitter release contain PI(4,5)P_2_-binding domains (Martin, 2001), PI(4,5)P_2_ is expected to be localized at the presynaptic active zones (AZs). The PI(4,5)P_2_ particle density was significantly higher in the AZs than in the whole bouton and extra-AZ membranes at both PF-PC and PF-MLI synapses (Tukey: p < 0.001; Figure 4B). The observed density of PI(4,5)P_2_ particles in AZs was significantly higher than that of randomly distributed PI(4,5)P_2_ particles on the bouton membrane by Monte-Carlo simulation (Chi-LRT: p < 0.001 (PF-PC) and p = 0.002 (PF-MLI); Figure 4C). Next, we examined how the PI(4,5)P_2_ particles are distributed within the AZs using the center-periphery index (CPI), where a CPI of 0 indicates that the particle is at the center of gravity, and a CPI of 1 indicates the particle is at the edge of the AZ (Kleindienst et al., 2020). The histogram of CPI in the bouton membranes suggests that the particles are distributed at a high density within the AZ and that the distribution probability decreases with distance from the AZ (Figure 4D, top). The CPI of the particles within AZs was higher at around CPI = 1.0 compared to the values of the random simulation (Figure 4D, bottom), indicating that PI(4,5)P_2_ is preferentially located in the periphery of the AZs. The mean CPI of the PI(4,5)P_2_ particles in AZs was significantly higher than that of randomly distributed particles (Chi-LRT: p < 0.001, Figure 4E) in both PF-PC and PF-MLI synapses. These results suggest that AZ, in particular, its periphery is PI(4,5)P_2_-enriched regardless of postsynaptic cell types.

**Figure 4.**
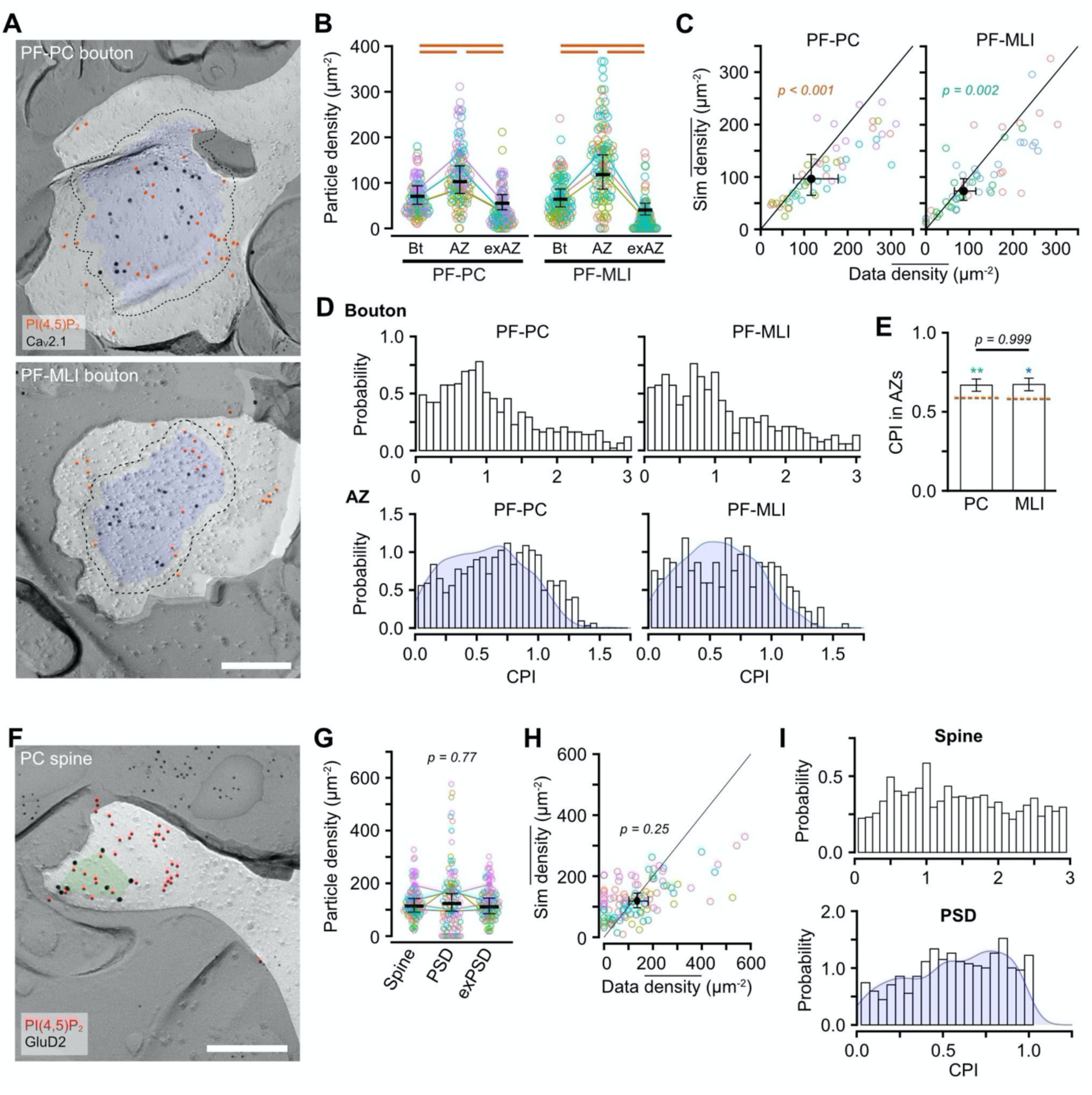
PI(4,5)P_2_ distribution in pre- and post-synaptic membranes of PF-PC synapses. (**A**) Example images for PI(4,5)P_2_ particle distribution on the membranes of PF-PC (top) and PF-MLI bouton (bottom). Red and black circles indicate gold particles for PI(4,5)P_2_ and Ca_V_2.1, respectively. The Blue area and dotted line indicate AZ and outer-rim (30 nm from the edge of AZ, Methods), respectively. Scale bar = 200 nm. (**B**) Beeswarm plot of the PI(4,5)P_2_ density in the whole bouton (Bt), AZs, and extra-AZ region (exAZ) of the PF-PC (left) and PF-MLI (right) bouton membranes. The PI(4,5)P_2_ density was significantly higher in AZs than in the whole bouton and the exAZ in both PF-PC and PF-MLI boutons (n = 111 boutons/4 mice, p < 0.001, Tukey method). (**C**) Comparison of the PI(4,5)P_2_ density in the AZs between real and simulated random distribution on the PF bouton membranes. The density of the real particle distribution was significantly higher than that of the simulated one in both PF-PC (real: 97.2 ± 19.7 particles/µm^2^, sim: 66.7 ± 13.5 particles/µm^2^, n = 55 boutons/4 mice, p < 0.001, Chi-LRT) and PF-MLI (real 86.7 ± 12.4 particles/µm^2^, sim: 73.2 ± 10.4 particles/µm^2^, n = 56 boutons/4 mice, p = 0.002, Chi-LRT) AZ membranes. (**D**) Distribution of center-periphery index (CPI) of PI(4,5)P_2_ particles in boutons (top) and AZs (bottom). Blue in the bottom graph indicates the CPI distribution of the simulated particles that are randomly distributed in AZs. (**E**) Comparison of CPIs of the PI(4,5)P_2_ particles in AZs between PF-PC and PF-MLI AZs. There is no significant difference in the CPIs between PF-PC and PF-MLI AZs (PF-PC: 0.67 ± 0.02, PF-MLI: 0.67 ± 0.02, n = 107 boutons/4 mice, p = 0.999, Chi-LRT). Dashed lines with red show the mean CPIs of the simulated randomly-distributed particles. Asterisks on the bars indicate statistical differences in CPIs between real and simulated particles (PF-PC: 0.59 ± 0.02, PF-MLI: 0.58 ± 0.02, *p < 0.05, **p < 0.01, Tukey method). (**F**) Example images of the PI(4,5)P_2_ particle distribution on the PC spine membrane. Red and black circles indicate PI(4,5)P_2_ and GluD2, respectively. The green area indicates postsynaptic density (PSD) based on the cluster of GluD2. Scale bar = 200 nm. (**G**) Beeswarm plot of the PI(4,5)P_2_ density in the whole spine (spine), PSD, and extra-PSD region (exPSD) of the PC spine membranes. No significant difference in the density was detected between these compartments (n = 108 spines/6 mice, p = 0.77, Chi-LRT). (**H**) Comparison of the PI(4,5)P_2_ density in the PSDs between real and simulated random distribution on the PC spine membranes. The density of the real and simulated particle distribution was not significantly different (real: 0.59 ± 0.01, sim: 0.55 ± 0.01, n = 108 spines/6 mice, p = 0.25, Chi-LRT). (**I**) Distribution of CPIs of PI(4,5)P_2_ particles in spines (top) and PSDs (bottom). Blue in the bottom graph indicates the CPI distribution of the simulated particles that are randomly distributed in PSDs. The CPI of the PI(4,5)P_2_ particles is uniformly distributed in the spines and PSDs, suggesting the random distribution of PI(4,5)P_2_.

Next, we investigated the PI(4,5)P_2_ distribution in PSD of dendritic spines. The PI(4,5)P_2_ particles were distributed throughout the spine membranes (Figure 4F), and the density in the PSDs was not significantly different compared to the whole spine and extra-PSD membranes (Chi-LRT: p = 0.77; Figure 4G). The observed PI(4,5)P_2_ particle density in PSDs was not significantly higher than that of the particles randomly distributed in the spine (Chi-LRT: p = 0.25; Figure 4H). Furthermore, the CPI of the PI(4,5)P_2_ particles in the spines was uniformly distributed, and the CPI in the PSDs was similar to the simulated particles randomly distributed (Figure 4I). These results suggest that PI(4,5)P_2_ is not specifically accumulated in the postsynaptic site of PC dendritic spines.

### The association of ion channels and receptors with PI(4,5)P_2_ on cell membranes of cerebellar neurons

#### The association of Ca_V_2.1 with PI(4,5)P_2_

Ca_V_2.1 is an α-subunit of P/Q-type voltage-gated calcium channels and is regulated by PI(4,5)P_2_ directly or indirectly through Ca_V_ β-subunit (Suh et al., 2012; Suh and Hille, 2008). Here, we examined whether Ca_V_2.1 is associated with PI(4,5)P_2_ in the neuronal membranes by co-immunolabeling of Ca_V_2.1 with the PI(4,5)P_2_ labeling. To assess the association, we compared NNDs from Ca_V_2.1 to PI(4,5)P_2_ particles (NND_C-P_) between observed and simulated Ca_V_2.1 particles. To reproduce the Ca_V_2.1 clustering in the simulation, we performed a fitted simulation of Ca_V_2.1 particle distribution (Methods, (Kleindienst et al., 2020; Luján et al., 2018b)). In PCs, Ca_V_2.1 is broadly expressed in somatodendritic membranes (Figure 5A) as previously reported (Indriati et al, 2013). The real values of NND_C-P_ in somatic, SpB, and spine membranes were significantly shorter than the simulated ones (Figure 5B). The mean NND_C-P_ was not significantly different between the compartments of PC membranes (Figure 5C). In GCs, Ca_V_2.1 was highly expressed in somatic and presynaptic AZ membranes (Figure 5D). The comparison of NND_C-P_ between observed and simulated Ca_V_2.1 particles showed significant differences in the somatic and PF-PC/MLI AZ membranes (Figure 5E), indicating the association of Ca_V_2.1 with PI(4,5)P_2_. The mean NND_C-P_ values in PF-PC and PF-MLI AZs were not significantly different (Figure 5F). In MLI, Ca_V_2.1 was expressed in somatic and presynaptic basket cell (BC)-PC bouton membranes (Figure 6A). The NND_C-P_ in somatic and BC-PC bouton membranes was significantly shorter than the simulated one (Figure 6B). These results suggest the ubiquitous association of Ca_V_2.1 with PI(4,5)P_2_ in neuronal cell membranes across various cell types and their compartments.

**Figure 5.**
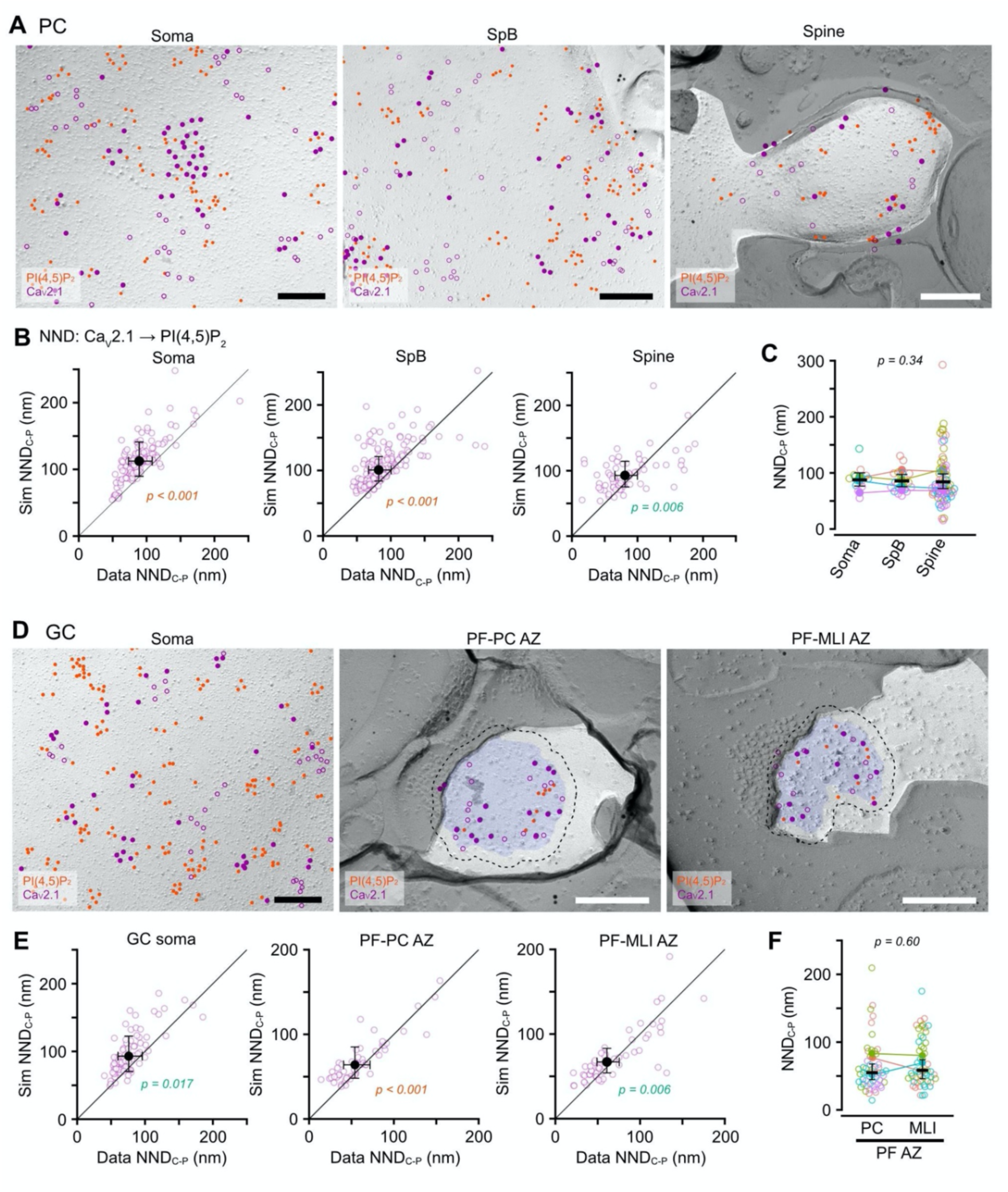
Ubiquitous association of Ca_V_2.1 with PI(4,5)P_2_ on cell membranes of PCs and GCs. (**A**) Example images for co-immunolabeling of PI(4,5)P_2_ and Ca_V_2.1 on somatic (left), SpB (middle), and spine (right) membranes of the PC. Red and purple (closed, open) circles indicate PI(4,5)P_2_ and Ca_V_2.1 (real, fitted-simulated) particles, respectively. Scale bars = 200 nm. (**B**) Comparison of the NNDs from Ca_V_2.1 to PI(4,5)P_2_ particles (NND_C-P_) between real and fitted-simulated Ca_V_2.1 distribution on somatic (left), SpB (middle), and spine (right) membranes of PCs. The NND_C-P_ of the real distribution was significantly smaller than that of the simulated one in somatic (real: 89.7 ± 9.6 nm, sim: 112.9 ± 12.1 nm, n = 129 images/14 cells/5 mice, p < 0.001, Chi-LRT), SpB (real: 82.3 ± 8.1 nm, sim: 100.5 ± 9.9 nm, n = 158 images/20 dendrites/4 mice, p < 0.001, Chi-LRT), and spine membrane (real: 80.9 ± 8.7 nm, sim: 92.8 ± 9.9 nm, 69 spines/4 mice, p = 0.006, Chi-LRT). (**C**) Comparison of NND_C-P_ between somatodendritic compartments. There is no significant difference between the PC compartments (n = 12,956 values/92 components/5 mice, p = 0.70, Chi-LRT). (**D**) Example images for co-immunolabeling of PI(4,5)P_2_ and Ca_V_2.1 on somatic (left) and presynaptic PF-PC (middle) and PF-MLI (right) AZ membranes of the GC. Red and purple (closed, open) circles indicate PI(4,5)P_2_ and Ca_V_2.1 (real, fitted-simulated) particles, respectively. The Blue area and dotted lines indicate AZs and the outer-rim, respectively. Scale bars = 200 nm. (**E**) Comparison of the NND_C-P_ between real and fitted-simulated Ca_V_2.1 distribution on somatic (left), PF-PC AZ (middle), and PF-MLI AZ (right) membranes of GCs.The NND_C-P_ of the real distribution was significantly smaller than that of the simulated one in somatic (real: 76.0 ± 9.0 nm, sim: 93.0 ± 13.1 nm, n = 81 images/25 cells/5 mice, p = 0.017, Chi-LRT), PF-PC AZ (real: 54.0 ± 7.9 nm, sim: 64.0 ± 9.3 nm, n = 54 AZs/4 mice, p < 0.001, Chi-LRT), and PF-MLI AZ membrane (real: 60.8 ± 6.7 nm, sim: 67.0 ± 7.3 nm, 52 AZs/4 mice, p = 0.006, Chi-LRT). (**F**) Comparison of NND_C-P_ between presynaptic AZs of PF-PC and PF-MLI synapses. There is no significant difference in NND_C-P_ between the AZs (n = 1,110 values/68 AZs/4 mice, p = 0.60, Chi-LRT).

**Figure 6.**
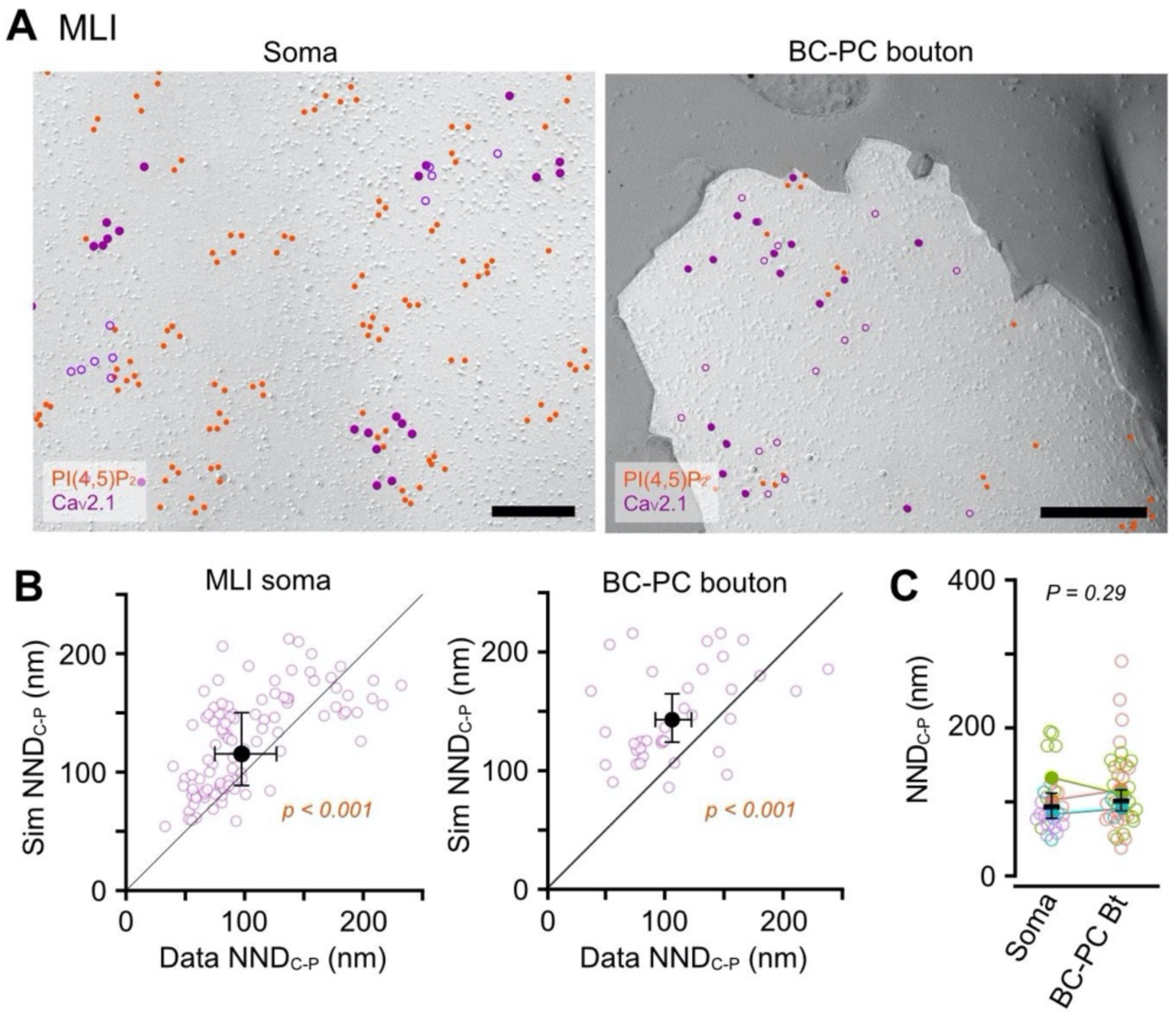
Association of Ca_V_2.1 with PI(4,5)P_2_ on cell membranes of MLIs. (**A**) Example images for co-immunolabeling of PI(4,5)P_2_ and Ca_V_2.1 on somatic (left) and basket cell (BC)-PC bouton (right) membranes of the MLI. Red and purple (closed, open) circles indicate PI(4,5)P_2_ and Ca_V_2.1 (real, fitted-simulated) particles, respectively. P-face of BC-PC boutons were identified based on Ca_V_2.1 clusters and the surrounding E-face of the PC somatic membranes with Ca_V_2.1 clusters. Scale bars = 200 nm. (**B**) Comparison of the NND_C-P_ between real and fitted-simulated Ca_V_2.1 distribution on somatic (left) and BC-PC bouton (right) membranes of MLIs. The NND_C-P_ of the real distribution was significantly smaller than that of the simulated one in somatic (real: 97.5 ± 13.1 nm, sim: 115.4 ± 15.5 nm, n = 102 images/26 cells/4 mice, p < 0.001, Chi-LRT) and BC-PC bouton membrane (real: 106.0 ± 7.7 nm, sim: 143.0 ± 10.4 nm, 37 boutons/3 mice, p = 0.006, Chi-LRT). (**C**) Comparison of NND_C-P_ between somatic and bouton membranes. No significant difference was shown between these compartments (n = 2,854 values/63 components/4 mice, p = 0.29, Chi-LRT).

#### The association of GIRK3 channels with PI(4,5)P_2_

PI(4,5)P_2_ is associated with some ion channels and regulates their activities, e.g., open channel probability (Suh and Hille, 2008). G protein-coupled inwardly rectifying K^+^ (GIRK) channels are a family of lipid-gated potassium channels that are activated by PI(4,5)P_2_ and G-protein βγ-subunits (G_βγ_) released from G-protein coupled receptors (GPCRs) (Whorton and MacKinnon, 2011). Thus, GIRK channels are expected to be associated with PI(4,5)P_2_ clusters in order to be efficiently activated. To examine whether PI(4,5)P_2_ and GIRK channels are co-clustered on dendritic membranes of mouse cerebellar PCs, we co-labeled PI(4,5)P_2_ and GIRK3, the most predominant subunit of GIRK channels in PCs (Aguado et al., 2008; Fernández-Alacid et al., 2009), on the P-face of the dendritic membranes. The GIRK3 particles were observed throughout the dendritic SpBs and spines of the PCs (Figure 7A). We compared the NNDs from GIRK3 to PI(4,5)P_2_ particles (NND_G-P_) between observed and simulated GIRK3 particles in spines and SpBs spines of PCs. The real NND_G-P_ in both spines and SpBs was significantly shorter than the simulated one (Chi-LRT: p < 0.001, Figure 7B), suggesting that GIRK3 channels co-clustered with PI(4,5)P_2_ in the distal dendritic membranes of PCs. GIRK channels are also expressed on presynaptic PF-PC boutons, including AZs (Figure 7A, (Fernández-Alacid et al., 2009; Luján et al., 2018b), and may regulate presynaptic excitability. In the presynaptic AZs, the real NND_G-P_ was significantly shorter than the simulated one (Chi-LRT: p < 0.001, Figure 7B), indicating the coexistence of GIRK3 with PI(4,5)P_2_. The mean NND_G-P_ in spine and AZ membranes was significantly shorter than that in SpB membranes (Figure 7C), suggesting the tighter association of GIRK3 and PI(4,5)P_2_ in synaptic membranes.

**Figure 7.**
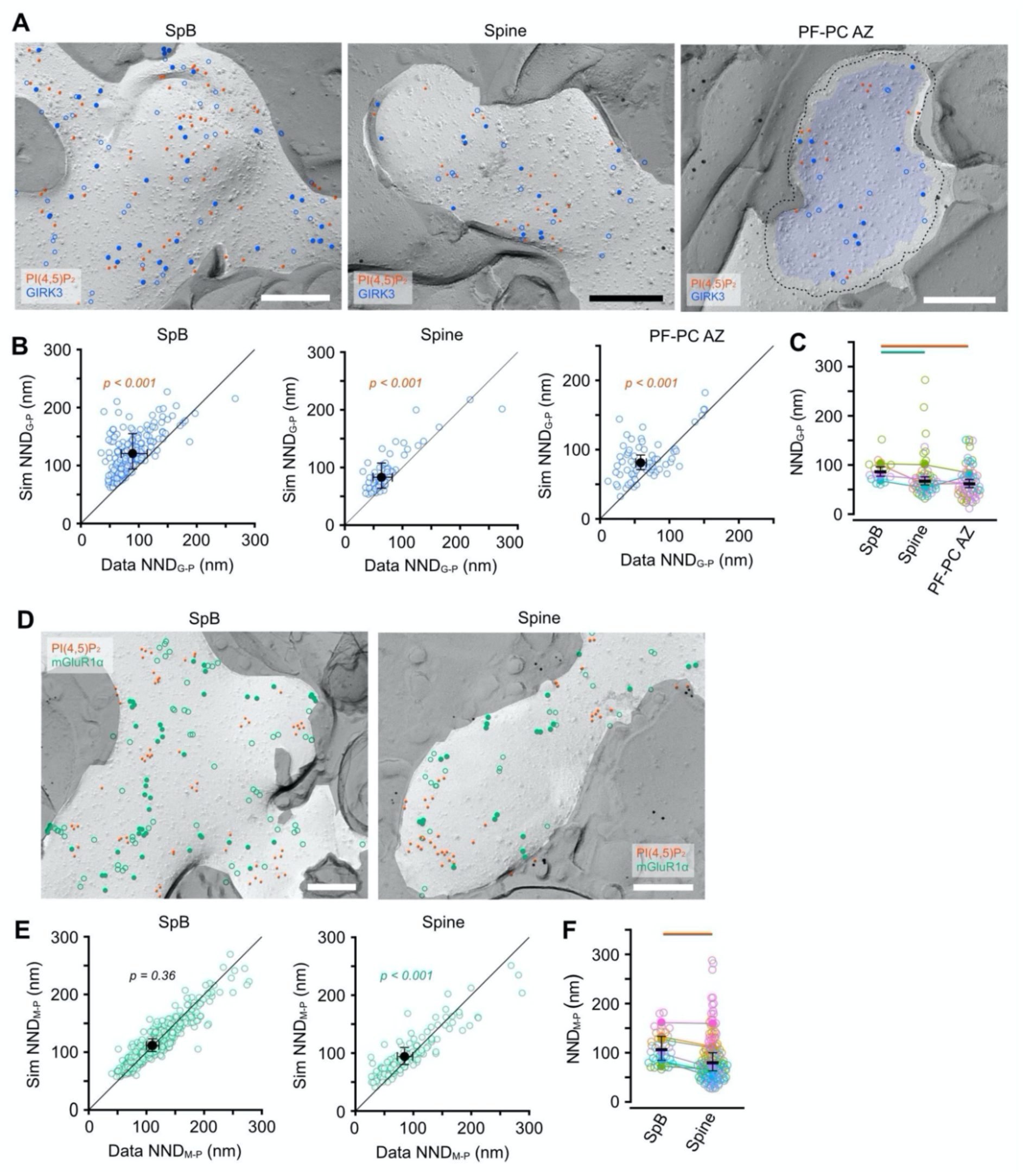
Association of GIRK3 channels and mGluR1α receptors with PI(4,5)P_2_ on cell membranes of cerebellar neurons. (**A**) Example images for co-labeling of PI(4,5)P_2_ and GIRK3 channels on PC SpB (left), PC spine (middle), and PF-PC AZ (right) membranes. Red and blue (closed, open) circles indicate PI(4,5)P_2_ and GIRK3 (real, fitted-simulated) particles, respectively. The Blue area and a dotted line on the left indicate the AZ and the outer-rim, respectively. Scale bars = 200 nm. (**B**) Comparison of the NNDs from GIRK3 to PI(4,5)P_2_ particles (NND_G-P_) between real and fitted-simulated GIRK3 distribution on PC SpB (left), PC spine (middle), and PF-PC AZ (right) membranes. The NND_G-P_ of the real distribution was significantly smaller than that of the simulated one in SpB (real: 89.3 ± 11.4 nm, sim: 120.7 ± 15.4 nm, n = 88 images/19 dendrites/5 mice, p < 0.001, Chi-LRT), spine (real: 63.6 ± 8.4 nm, sim: 83.0 ± 11.0 nm, n = 57 spines/4 mice, p < 0.001, Chi-LRT), and PF-PC AZ membrane (real: 58.6 ± 3.9 nm, sim: 81.1 ± 5.3 nm, 65 AZs/4 mice, p < 0.001, Chi-LRT). (**C**) Comparison of NND_G-P_ between the post- and presynaptic compartments of cerebellar neurons. The NND_G-P_ was significantly shorter in the spine and AZ membranes than in the SpB membrane (4,730 values/90 compartments/4 mice, p < 0.001, Chi-LRT). (**D**) Example images for co-labeling of PI(4,5)P_2_ and mGluR1α receptors on PC SpB (left) and spine (right) membranes. Red and green (closed, open) circles indicate PI(4,5)P_2_ and mGluR1α (real, fitted-simulated) particles, respectively. Scale bars = 200 nm. (**E**) Comparison of the NNDs from mGluR1α to PI(4,5)P_2_ particles (NND_M-P_) between real and fitted-simulated mGluR1α distribution on PC SpB (left) and spine (right) membranes. The NND_M-P_ of the real distribution was significantly smaller than that of the simulated one in the spine membranes (real: 81.8 ± 18.4 nm, sim: 91.2 ± 20.6 nm, n = 140 spines/7 mice, p < 0.001, Chi-LRT), but not in the SpB membrane (n = 147 images/35 dendrites/7 mice, p = 0.36, Chi-LRT). (**F**) Comparison of NND_M-P_ between the SpB and spine membranes of PCs. The NND_M-P_ was shorter in the spines than in the SpB membrane (real: 110.0 ± 5.6 nm, sim: 112.0 ± 5.6 nm, n = 22,391 values/140 compartments/7 mice, p < 0.001, Chi-LRT).

#### The association of mGluR1α with PI(4,5)P_2_

Group I metabotropic glutamate receptors (mGluRs) are GPCRs coupled with Gα_q_ subunit and hydrolyze PI(4,5)P_2_ into inositol 1,4,5-trisphosphate (IP_3_) and diacylglycerol (DAG) through the activation of PLCβ. A subtype of Group I mGluRs mGluR1α is highly expressed on the dendritic shafts and spines but avoids PSDs in PCs, and forms clusters (Luján et al., 2018a). To effectively produce the second messengers from PI(4,5)P_2_, a mGluR1α cluster is expected to be located close to the PI(4,5)P_2_ clusters on the dendritic membrane of PCs. To address this possibility, we co-labeled mGluR1α with PI(4,5)P_2_ on the dendritic membranes of PCs and compared the NNDs between mGluR1α and PI(4,5)P_2_ particles (NND_M-P_) with the fitted simulation of the mGluR1α particle distribution. The mGluR1α particles were distributed as clusters on the membranes of spines and SpBs (Figure 7D-E) as previously reported (Luján et al., 2018a). The real NND_M-P_ was significantly shorter than the simulated one on the spine membrane (Chi-LRT: p < 0.001) but not on the SpB membrane (Chi-LRT: p = 0.36, Figure 7F). These results suggest compartment-specific co-clustering of mGluR1α with PI(4,5)P_2_ on PC spines.

## Discussion

### Visualization of PI(4,5)P_2_ distribution on neuronal cell membranes using SDS-FRL

To understand how and where PI(4,5)P_2_ exerts the physiological functions in neurons, it is essential to know its nanoscale distribution in the neuronal cell membranes. To visualize the distribution of PI(4,5)P_2_ on the cell membrane, cryofixation of phospholipids is necessary because they diffuse laterally even after chemical fixation (Tanaka et al., 2010). Detergents and organic solvents used in conventional immunoEM methods disrupt the cell membrane and should be avoided. In regions with densely packed proteins such as AZs and PSDs, limited accessibility of the probes could hamper quantitative analysis of the PI(4,5)P_2_ distribution. The fluorescent protein-tagged PH domain of PLCδ1 has been developed as a specific probe for PI(4,5)P_2_ (Idevall-Hagren and De Camilli, 2015). This probe allows us to perform single-molecule imaging of PI(4,5)P_2_ on the plasma membrane of the cultured cells using super-resolution microscopies such as Stimulated Emission Depletion (STED) microscopy and Direct Stochastic Optical Reconstruction Microscopy (dSTORM) (Milovanovic et al., 2016; Wang and Richards, 2012). However, it is still difficult to observe the PI(4,5)P_2_ distribution on neuronal cell membranes in brain tissues because of the limited accessibility of the probes in AZs and PSDs. In addition, overexpressed fluorescent probes interfere with endogenous protein binding to PI(4,5)P_2_ by masking PI(4,5)P_2_ on the cell membrane (Suh and Hille, 2008), making it difficult to quantify the changes in PI(4,5)P_2_. Thus, the SDS-FRL with cryo-fixed brain tissue used in this study has the advantage of visualizing accurately the native PI(4,5)P_2_ distribution. In this method, phospholipids are physically immobilized by high-pressure freezing and replication by deposition of evaporated carbon/platinum, so the phospholipids do not diffuse after freezing. Since cytosolic proteins are removed by SDS treatment, PI(4,5)P_2_ and transmembrane proteins are exposed on the replica surface, making high accessibility of the probes and antibodies. Due to a low temporal resolution, it is still challenging to examine the dynamics of PI(4,5)P_2_ during neuronal activity, but it is a beneficial method to study the nanoscale distribution of PI(4,5)P_2_ and other phospholipids on neuronal cell membranes.

This study demonstrates that PI(4,5)P_2_ is broadly distributed on neuronal cell membranes in the mouse cerebellum. Comparisons of the PI(4,5)P_2_ distribution between cell types revealed slightly different densities of PI(4,5)P_2_ particles on the somatic membrane, but the differences in distribution patterns between subcellular compartments were more pronounced; in PCs, the PI(4,5)P_2_ density was higher at the distal dendrites than at the somatic and proximal dendrites; in GCs, the density was similar at the somata, dendrites, axons, and presynaptic boutons; in MLIs, the density of PI(4,5)P_2_ in clusters was lower in presynaptic bouton membranes than somatic and dendritic membranes. These differences in the PI(4,5)P_2_ distribution patterns may reflect different roles of PI(4,5)P_2_ in cell functions between different cell types.

### PI(4,5)P_2_ makes clusters on neuronal cell membranes in mouse cerebellum

Previous studies using super-resolution microscopy have demonstrated that PI(4,5)P_2_ forms clusters of 50-100 nm in diameter on the cell membrane, as shown in PC12 cells (Aoyagi et al., 2005; van den Bogaart et al., 2011; Wang and Richards, 2012) and mouse myoblast cells (Petersen et al., 2016). Our study reveals that PI(4,5)P_2_ forms clusters of around 1,000 nm^2^(corresponding to 35 nm in diameter) on the cell membrane of neurons in the acute cerebellar tissues. The area of the clusters showed almost the same mean values in the somatodendritic and axonal membranes of different neuronal cell types in the cerebellum. The cluster density and the intra-cluster particle density of PI(4,5)P_2_ in the PC spine membrane are higher than in other somatodendritic compartments. These observations suggest the accumulation of densely-packed PI(4,5)P_2_ clusters in the spine membranes, supporting the contribution of PI(4,5)P_2_ and other PIs to spine formation and morphological long-term plasticity (Lei et al., 2017; Ueda and Hayashi, 2013). Although it is still unknown why PI(4,5)P_2_ forms clusters, the clustered PI(4,5)P_2_ localization indicates the importance of a close spatial relationship between the clusters and effector proteins for their effective interaction, as discussed in the following section.

### PI(4,5)P_2_ distribution in the synaptic membrane

In this study, we found the accumulation of PI(4,5)P_2_ in AZ membranes of the presynaptic PF boutons regardless of the postsynaptic cell types. This result suggests the roles of PI(4,5)P_2_ on presynaptic activities, including vesicle exocytosis and endocytosis. At the presynaptic membrane, Rab6-interacting proteins (RIMs) have PI(4,5)P_2_-binding sites at C2A and C2B domains, and their interactions are essential for efficient exocytosis (de Jong et al., 2018). PI(4,5)P_2_ also interacts with a SNARE protein syntaxin-1A and an SV Ca^2+^ sensor protein synaptotagmin-1 (Aoyagi et al., 2005; Honigmann et al., 2013; Milovanovic et al., 2016). Since Ca_V_2.1 directly binds to RIMs and dominantly induces Ca^2+^ influx into PF boutons, the association of Ca_V_2.1 with PI(4,5)P_2_ supports the idea that PI(4,5)P_2_ anchors SVs and Ca^2+^ channels through interaction with these proteins. PI(4,5)P_2_ also regulates vesicle endocytosis via interactions with endocytic proteins e.g., AP-2, AP180, clathrin, and dynamin (Haucke, 2005; Martin, 2001; Posor et al., 2015). At calyx of Held synapses of rat brainstem, upregulation of PI(4,5)P_2_ by the retrograde signals through the nitric oxide/cyclic GMP-activated protein kinase/RhoA/Rho-associated protein kinase pathway accelerates vesicle endocytosis, which strengthens the homeostatic plasticity for the maintenance of high-frequency synaptic transmission (Eguchi et al., 2012; Taoufiq et al., 2013). PI(4,5)P_2_ clusters may work as anchors of presynaptic proteins involved in this pathway providing a site for their interactions.

### Co-clustering of PI(4,5)P_2_ and ion channels and receptors

PI(4,5)P_2_ regulates ion channels such as inwardly rectifying K^+^ channels, KCNQ channels, transient receptor potential (TRP) channels, and voltage-gated Ca^2+^ channels (Suh and Hille, 2008). Therefore, clarifying the distance relationship between these ion channels and PI(4,5)P_2_ may provide insight into the regulation mechanism of neuronal excitability by PI(4,5)P_2_. In this study, we found that GIRK3 was associated with PI(4,5)P_2_ in SpB and spines of PCs and also AZs of PF boutons. Because PI(4,5)P_2_ is essential to activate GIRK channels by βγ subunit of G protein (Whorton and MacKinnon, 2011), the association between GIRK3 and PI(4,5)P_2_ suggests that GIRK3 on SpB and spines is constantly ready to be activated by G-protein coupled receptors. Since GIRK channels are also associated with GABA_B_ receptors at different compartments of the cerebellar neurons, such as dendritic shafts and spines of PCs and AZs of PF boutons (Fernández-Alacid et al., 2009; Luján et al., 2018b), the co-assembly of PI(4,5)P_2_, GIRK channels and GABA_B_ receptors may contribute to effective inhibitory postsynaptic transmission on Purkinje cells (Luján et al., 2018b). Group I mGluRs are GPCRs coupled with Gα_q_ subunit and hydrolyze PI(4,5)P_2_ into inositol 1,4,5-trisphosphate (IP_3_) and diacylglycerol (DAG) through the activation of PLCβ. PCs express mGluR1α, a subtype of Group I mGluRs, on the dendritic shafts and spines concentrated in the peri-PSD areas (Luján et al., 2018a). We found a significant association of mGluR1α with PI(4,5)P_2_ clusters specific to the spine membrane in PCs, which may contribute to the effective production of the second messengers from PI(4,5)P_2_ in response to the glutamate release from presynaptic PF boutons. On the other hand, no particular concentration of PI(4,5)P_2_ has been found in the peri-PSD areas. The diffusion dynamics of PI(4,5)P_2_ on the spine membrane may be enough to support the interaction with mGluR1α. Thus, the tight spatial association of PI(4,5)P_2_ as observed for Ca_V_2.1 and GIRK3 may not be necessary for the coupling of mGluR1α activation and intracellular signal cascade, though the distance between mGluR1α (and the coupled PLCβ) and PI(4,5)P_2_ may affect the kinetics of the second messenger (IP_3_ and DAG) production from PI(4,5)P_2_.

The present study demonstrated that the SDS-FRL method could be successfully applied to neuronal cells in acute mouse brain tissues to visualize the distribution of PI(4,5)P_2_. We quantitatively analyzed the nanoscale two-dimensional distribution of PI(4,5)P_2_ on the cell membrane of mouse cerebellar neurons, and its spatial relationship with GIRK3, Ca_V_2.1, and mGluR1α in PC dendrites and PF boutons. Notably, we have demonstrated a higher density of PI(4,5)P_2_ in distal PC dendrites, showing a specific association of mGluR1α with PI(4,5)P_2_ in spines. PI(4,5)P_2_ also showed concentration and association with Ca_V_2.1 and GIRK3 in presynaptic PF AZs. These results will help to elucidate the physiological functions of PI(4,5)P_2_ in neuronal activities, including SV fusion and retrieval in presynaptic membranes and long-term plasticity in postsynaptic spine membranes. This method could also be a powerful tool for the analysis of other PIs and phospholipid distribution in neuronal cell membranes using specific probes.

## Author Contributions

KE and RS designed and conducted the experiments and wrote the manuscript. KE analyzed the data.

## Acknowledgments

We would like to thank Nicoleta Condruz (IST Austria, Klosterneuburg, Austria) for technical assistance with sample preparation; the Electron Microscopy Facility of IST Austria (Klosterneuburg, Austria) for technical support with EM works; Natalia Baranova (University of Vienna, Vienna, Austria) and Martin Loose (IST Austria, Klosterneuburg, Austria) for advice on liposome preparation; Yugo Fukazawa (University of Fukui, Fukui, Japan) for comments. This work was financially supported by funding from the Institute of Science and Technology Austria, the European Union’s Horizon 2020 research and innovation program under the Marie Skłodowska-Curie grant agreement No. 793482 (to KE), and the European Research Council (ERC) grant agreement No. 694539 (to RS).

