## Supplemental Information for "Nanoscale phosphoinositide distribution on cell membranes of mouse cerebellar neurons"

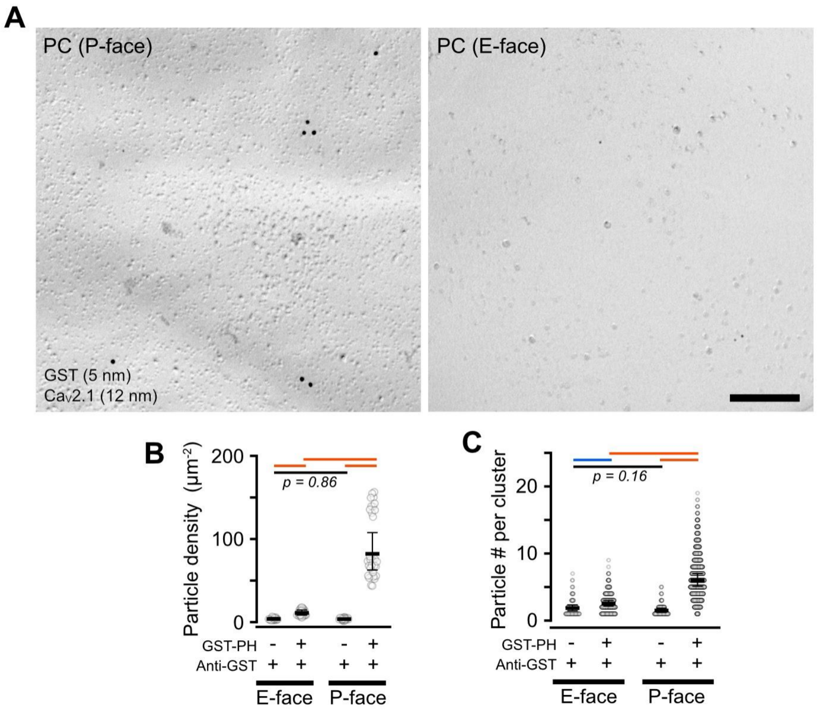


Figure S1. Non-specific labeling on the somatic membranes of PCs. Related to Figure 1.

(**A**) Example images of the $PI(4,5)P_2$ labeling without incubation with GST-PH on the P- (left) and E-face of PC somatic membranes. The replicas were subsequently incubated with anti-GST/anti-$Ca_V2.1$ primary antibodies and gold-conjugated secondary antibodies (5 nm for anti-GST, 12 nm for anti-$Ca_V2.1$). Scale bar = 200 nm.

(**B**) Comparison of the 5-nm gold particle densities with and without GST-PH on the E- and P-face of PC somatic membranes. The particle density without GST-PH was significantly lower than that with GST-PH in both faces. Without the GST-PH incubation, no significant difference in the density was detected between the E- and P-face ($p = 0.86$, multiple comparisons with the Benjamini-Hochberg (BH) method).

(**C**) Comparison of the number of 5-nm gold particles per cluster on the E- and P-face of the PC somatic membranes incubated with or without GST-PH. The clusters were detected using Ward Linkage hierarchical clustering method. The threshold distance to separate each cluster was set as 50 nm, which is nearby the maximum distance for cluster detection on the PC somatic membrane using DBSCAN (Figure 2, Method). The particle number per cluster without GST-PH incubation was significantly lower than that with GST-PH in both faces (multiple comparisons with BH method), suggesting that the $PI(4,5)P_2$ particle clusters observed on the neuronal membranes are also ascribable to specific clustering of GST-PH binding sites but not to non-specific aggregation of the primary and secondary antibodies.

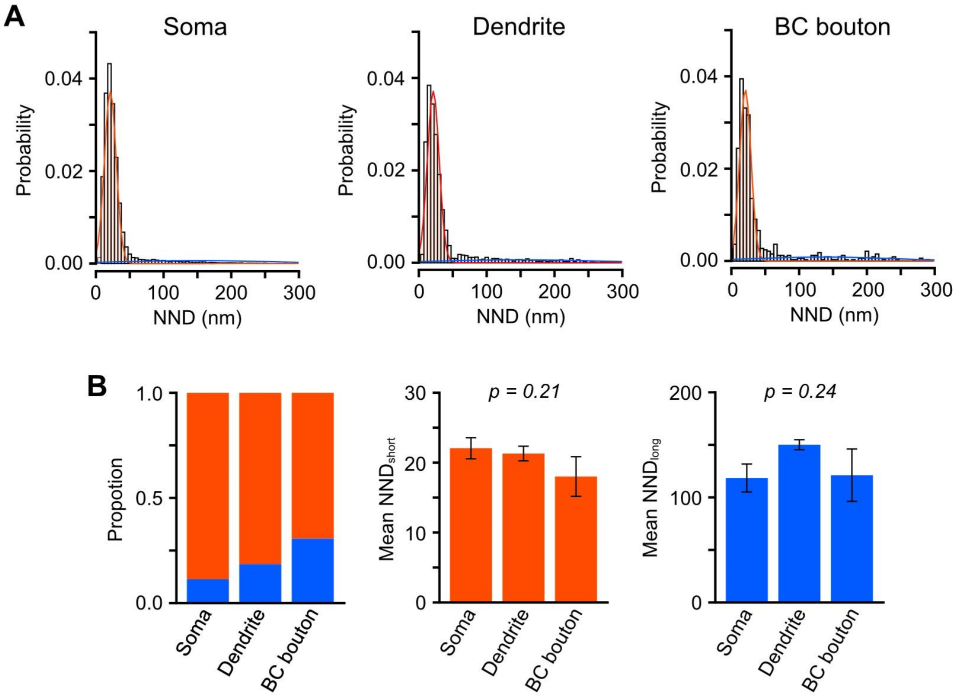


**Figure S2. Clustered and scattered PI(4,5)P$_2$ distribution on MLI compartments. Related to Figure 3.**

(**A**) Distribution of NNDs distribution of the PI(4,5)P$_2$ particles obtained from the somatic (left, n = 20,164 particles), dendritic (middle, n = 3,175 particles), and BC bouton membranes (right, n = 673 particles) of MLIs. Red and blue lines indicate the shorter and longer components of the NND distribution, respectively, estimated from the Gaussian mixture modeling.

(**B**) Proportion (left) and mean values of shorter (red, middle) and longer (blue, left) components of the NND of PI(4,5)P$_2$ particles on MLI compartments estimated by the Gaussian mixture model. Since there are no significant differences in the mean values of both shorter and longer components of the NND ($p = 0.21$ and 0.24, respectively, Chi-LRT), the higher mean NND values in the dendritic and BC bouton membranes compared to the somatic membranes (Figure 3E) are ascribable to larger proportions of the scattered PI(4,5)P$_2$ particles in these compartments.

**Table S1. PI(4,5)$P_2$ distribution in somatodendritic membrane compartments of PCs. Related to Figure 2.**

*Particle density ($\mu m^{-2}$)*

| Compartment* | mean** | s.e.m | 95% CI | | P-value† / Z-score | | | | |
|---|---|---|---|---|---|---|---|---|---|
| | | | Lower | Upper | Spine | SpB | SmB | MS | Soma |
| Soma | 52.3 | 7.0 | 40.2 | 68.1 | < .001 | 0.341 | 0.949 | 1.000 | - |
| MS | 52.3 | 7.1 | 40.2 | 68.2 | < .001 | 0.346 | 0.949 | - | -0.002 |
| SmB | 48.7 | 6.0 | 38.2 | 62.1 | < .001 | 0.019 | - | 0.732 | 0.732 |
| SpB | 62.7 | 7.8 | 49.2 | 80.0 | 0.149 | - | -3.059 | -1.848 | -1.858 |
| Spine | 73.7 | 8.6 | 58.6 | 92.5 | - | -2.287 | -5.882 | -3.917 | -3.917 |

*NND (nm)*

| Compartment* | mean** | s.e.m | 95% CI | | P-value† / Z-score | | | | |
|---|---|---|---|---|---|---|---|---|---|
| | | | Lower | Upper | Spine | SpB | SmB | MS | Soma |
| Soma | 34.3 | 2.3 | 30.2 | 39.0 | 0.877 | 1.000 | 0.871 | 0.992 | - |
| MS | 33.5 | 2.3 | 29.3 | 38.4 | 0.983 | 0.994 | 0.662 | - | 0.444 |
| SmB | 35.9 | 2.3 | 31.6 | 40.8 | 0.419 | 0.755 | - | -1.347 | -0.964 |
| SpB | 34.2 | 2.1 | 30.3 | 38.6 | 0.873 | - | 1.195 | -0.416 | 0.089 |
| Spine | 32.4 | 2.4 | 28.0 | 37.6 | - | 0.960 | 1.724 | 0.544 | 0.950 |

*Cluster density ($\mu m^{-2}$)*

| Compartment* | mean** | s.e.m | 95% CI | | P-value† / Z-score | | | | |
|---|---|---|---|---|---|---|---|---|---|
| | | | Lower | Upper | Spine | SpB | SmB | MS | Soma |
| Soma | 8.28 | 1.00 | 6.55 | 10.5 | < .001 | 0.483 | 0.859 | 0.990 | - |
| MS | 7.95 | 0.96 | 6.27 | 10.1 | < .001 | 0.221 | 0.991 | - | 0.467 |
| SmB | 7.67 | 0.88 | 6.13 | 9.60 | < .001 | 0.027 | - | 0.455 | 0.993 |
| SpB | 9.41 | 1.08 | 7.51 | 11.8 | 0.054 | - | -2.943 | -2.098 | -1.622 |
| Spine | 11.7 | 1.42 | 9.22 | 14.8 | - | -2.701 | -5.321 | -4.349 | -3.945 |

*Cluster area ($nm^2$)*

| Compartment* | mean** | s.e.m | 95% CI | | P-value† / Z-score | | | | |
|---|---|---|---|---|---|---|---|---|---|
| | | | Lower | Upper | Spine | SpB | SmB | MS | Soma |
| Soma | 936 | 76 | 798 | 1098 | 0.265 | 0.960 | 0.999 | 1.000 | - |
| MS | 914 | 88 | 757 | 1105 | 0.385 | 0.927 | 0.991 | - | 0.185 |
| SmB | 965 | 64 | 847 | 1099 | 0.155 | 0.992 | - | -0.457 | -0.288 |
| SpB | 1006 | 68 | 882 | 1148 | 0.085 | - | -0.443 | -0.812 | -0.684 |
| Spine | 668 | 99 | 500 | 892 | - | 2.524 | 2.270 | 1.780 | 2.003 |

*Intra-cluster particle density ($nm^{-2}$)*

| Compartment* | mean** | s.e.m | 95% CI | | P-value† / Z-score | | | | |
|---|---|---|---|---|---|---|---|---|---|
| | | | Lower | Upper | Spine | SpB | SmB | MS | Soma |
| Soma | 0.0093 | 0.0007 | 0.0080 | 0.0108 | < .001 | 0.244 | 0.996 | 0.910 | - |
| MS | 0.0100 | 0.0008 | 0.0086 | 0.0117 | 0.006 | 0.826 | 0.977 | - | -0.865 |
| SmB | 0.0096 | 0.0007 | 0.0084 | 0.0109 | < .001 | 0.319 | - | 0.588 | -0.376 |
| SpB | 0.0109 | 0.0008 | 0.0095 | 0.0125 | 0.055 | - | -1.896 | -1.063 | -2.047 |
| Spine | 0.0137 | 0.0011 | 0.0117 | 0.0160 | - | -2.693 | -4.370 | -3.394 | -4.308 |

*MS = main shaft, SmB = smooth branchlet, SpB = spiny branchlet

**Marginal means estimated from generalized mixed-effects models

†P-values are calculated using mutiple comparison with Tukey method

**Table S2. $PI(4,5)P_2$ distribution in different membrane compartments of GCs. Related to Figure 3.**

*Particle density ($\mu m^{-2}$)*

| | | | 95% CI | | P-value† / Z-score | | | | |
|---|---|---|---|---|---|---|---|---|---|
| **Compartment*** | **mean**** | **s.e.m** | **Lower** | **Upper** | **PF-MLI** | **PF-PC** | **PF axon** | **Dendrite** | **Soma** |
| Soma | 68.8 | 10.8 | 51.6 | 94.4 | 0.961 | 1.000 | 0.339 | 0.658 | - |
| Dendrite | 57.7 | 7.2 | 45.3 | 73.6 | 0.916 | 0.309 | 0.916 | - | 1.353 |
| PF axon | 53.2 | 6.9 | 41.2 | 68.7 | 0.540 | 0.088 | - | 0.845 | 1.860 |
| PF-PC bouton | 70.9 | 8.9 | 55.3 | 90.7 | 0.864 | - | -2.510 | -1.916 | -0.108 |
| PF-MLI bouton | 63.0 | 8.5 | 48.4 | 82.1 | - | 0.980 | -1.534 | -0.847 | 0.679 |

*NND (nm)*

| | | | 95% CI | | P-value† / Z-score | | | | |
|---|---|---|---|---|---|---|---|---|---|
| **Compartment*** | **mean**** | **s.e.m** | **Lower** | **Upper** | **PF-MLI** | **PF-PC** | **PF axon** | **Dendrite** | **Soma** |
| Soma | 30.8 | 1.8 | 27.6 | 34.5 | 0.153 | 0.054 | 0.027 | 0.521 | - |
| Dendrite | 34.5 | 2.4 | 30.2 | 39.4 | 0.889 | 0.848 | 0.298 | - | -1.563 |
| PF axon | 43.7 | 5.1 | 34.8 | 54.9 | 0.775 | 0.726 | - | -1.937 | -2.944 |
| PF-PC bouton | 37.4 | 2.6 | 32.7 | 42.8 | 1.000 | - | 1.243 | -1.017 | -2.701 |
| PF-MLI bouton | 37.5 | 3.1 | 31.9 | 44.2 | - | -0.023 | 1.158 | -0.922 | -2.274 |

*Cluster density ($\mu m^{-2}$)*

| | | | 95% CI | | P-value† / Z-score | | | | |
|---|---|---|---|---|---|---|---|---|---|
| **Compartment*** | **mean**** | **s.e.m** | **Lower** | **Upper** | **PF-MLI** | **PF-PC** | **PF axon** | **Dendrite** | **Soma** |
| Soma | 10.94 | 1.51 | 8.35 | 14.3 | 0.887 | 0.994 | 0.357 | 0.541 | - |
| Dendrite | 9.11 | 1.20 | 7.04 | 11.8 | 0.997 | 0.832 | 0.937 | - | 1.534 |
| PF axon | 8.14 | 1.39 | 5.82 | 11.4 | 0.885 | 0.582 | - | 0.780 | 1.828 |
| PF-PC bouton | 10.37 | 1.47 | 7.86 | 13.7 | 0.982 | - | -1.470 | -1.050 | 0.414 |
| PF-MLI bouton | 9.5 | 1.51 | 6.99 | 13.0 | - | 0.552 | -0.931 | -0.347 | 0.926 |

*Cluster area ($nm^2$)*

| | | | 95% CI | | P-value† / Z-score | | | | |
|---|---|---|---|---|---|---|---|---|---|
| **Compartment*** | **mean**** | **s.e.m** | **Lower** | **Upper** | **PF-MLI** | **PF-PC** | **PF axon** | **Dendrite** | **Soma** |
| Soma | 1016 | 144 | 770 | 1340 | 0.990 | 0.971 | 0.772 | 0.053 | - |
| Dendrite | 696 | 127 | 488 | 995 | 0.974 | 0.340 | 1.000 | - | 2.705 |
| PF axon | 673 | 215 | 360 | 1260 | 0.992 | 0.749 | - | 0.091 | 1.164 |
| PF-PC bouton | 1208 | 442 | 590 | 2475 | 0.869 | - | -1.205 | -1.859 | -0.924 |
| PF-MLI bouton | 851 | 327 | 400 | 1808 | - | 0.969 | -0.439 | -0.610 | 0.558 |

*Intra-cluster particle density ($nm^{-2}$)*

| | | | 95% CI | | P-value† / Z-score | | | | |
|---|---|---|---|---|---|---|---|---|---|
| **Compartment*** | **mean**** | **s.e.m** | **Lower** | **Upper** | **PF-MLI** | **PF-PC** | **PF axon** | **Dendrite** | **Soma** |
| Soma | 0.0092 | 0.0009 | 0.0077 | 0.0110 | 0.989 | 0.991 | 0.993 | 0.119 | - |
| Dendrite | 0.0123 | 0.0017 | 0.0094 | 0.0160 | 0.845 | 0.370 | 0.703 | - | -2.385 |
| PF axon | 0.0082 | 0.0022 | 0.0048 | 0.0139 | 0.976 | 1.000 | - | 1.281 | 0.432 |
| PF-PC bouton | 0.0086 | 0.0014 | 0.0063 | 0.0117 | 0.968 | - | -0.163 | 1.807 | 0.455 |
| PF-MLI bouton | 0.0101 | 0.0023 | 0.0065 | 0.0157 | - | -0.642 | -0.593 | 1.023 | -0.480 |

*PF= parallel fiber, PC = Purkinje cell, MLI = molecular layer interneuron

**Marginal means estimated from generalized mixed-effects models

†P-values are calculated using mutiple comparison with Tukey method

### Table S3. PI(4,5)$P_2$ distribution in different membrane compartments of MLIs

***Particle density ($\mu m^{-2}$)***

| | | | 95% CI | | P-value† / Z-score | | |
|---|---|---|---|---|---|---|---|
| Compartment* | mean** | s.e.m | Lower | Upper | BC-PC | Dendrite | Soma |
| Soma | 42.4 | 6.5 | 31.4 | 57.1 | 0.449 | 0.789 | - |
| Dendrite | 39.1 | 5.6 | 29.5 | 51.9 | 0.649 | - | 0.656 |
| BC-PC bouton | 35.7 | 5.8 | 26.1 | 49.0 | - | 0.887 | 1.208 |

***NND (nm)***

| | | | 95% CI | | P-value† / Z-score | | |
|---|---|---|---|---|---|---|---|
| Compartment* | mean** | s.e.m | Lower | Upper | BC-PC | Dendrite | Soma |
| Soma | 34.9 | 2.6 | 30.3 | 40.3 | < .001 | < .001 | - |
| Dendrite | 44.9 | 3.5 | 38.5 | 52.2 | 0.358 | - | -3.612 |
| BC-PC bouton | 52.2 | 6.1 | 41.5 | 65.7 | - | -1.368 | -3.650 |

***Cluster density ($\mu m^{-2}$)***

| | | | 95% CI | | P-value† / Z-score | | |
|---|---|---|---|---|---|---|---|
| Compartment* | mean** | s.e.m | Lower | Upper | BC-PC | Dendrite | Soma |
| Soma | 6.73 | 1.08 | 4.91 | 9.21 | 0.417 | 0.635 | - |
| Dendrite | 6.02 | 0.96 | 4.40 | 8.24 | 0.791 | - | 0.908 |
| BC-PC bouton | 5.49 | 1.05 | 3.77 | 8.00 | - | 0.652 | 1.261 |

***Cluster area ($nm^2$)***

| | | | 95% CI | | P-value† / Z-score | | |
|---|---|---|---|---|---|---|---|
| Compartment* | mean** | s.e.m | Lower | Upper | BC-PC | Dendrite | Soma |
| Soma | 979 | 303 | 534 | 1794 | 0.994 | 0.307 | - |
| Dendrite | 690 | 115 | 498 | 955 | 0.667 | - | 1.468 |
| BC-PC bouton | 931 | 263 | 535 | 1619 | - | -0.857 | 0.109 |

***Intra-cluster particle density ($nm^{-2}$)***

| | | | 95% CI | | P-value† / Z-score | | |
|---|---|---|---|---|---|---|---|
| Compartment* | mean** | s.e.m | Lower | Upper | BC-PC | Dendrite | Soma |
| Soma | 0.0103 | 0.0011 | 0.0083 | 0.0127 | 0.008 | 0.747 | |
| Dendrite | 0.0111 | 0.0009 | 0.0095 | 0.0129 | 0.016 | | -0.727 |
| BC-PC bouton | 0.0038 | 0.0015 | 0.0017 | 0.0084 | | 2.763 | 2.975 |

*BC = basket cell

**Marginal means estimated from generalized mixed-effects models

†P-values are calculated using mutiple comparison with Tukey method

**Table S4. PI(4,5)$P_2$ particle density in synaptic membranes of cerebellar neurons (particles/$\mu m^2$). Related to Figure 4.**

***PF-PC boutons***

| | | | 95% CI | | P-value[†] / Z-score | | |
|---|---|---|---|---|---|---|---|
| **Compartment**[*] | **mean**[**] | **s.e.m** | **Lower** | **Upper** | **exAZ** | **AZ** | **Bouton** |
| Bouton | 70.2 | 10.2 | 52.9 | 93.3 | < .001 | < .001 | - |
| AZ | 102.6 | 15.2 | 76.7 | 137.2 | < .001 | - | -5.908 |
| exAZ | 55.1 | 8.3 | 40.9 | 74.1 | - | 6.952 | 4.018 |

***PF-MLI boutons***

| | | | 95% CI | | P-value[†] / Z-score | | |
|---|---|---|---|---|---|---|---|
| **Compartment**[*] | **mean**[**] | **s.e.m** | **Lower** | **Upper** | **exAZ** | **AZ** | **Bouton** |
| Bouton | 63.8 | 9.9 | 47.2 | 86.4 | < .001 | < .001 | - |
| AZ | 118.0 | 18.9 | 86.2 | 161.3 | < .001 | - | -8.089 |
| exAZ | 40.3 | 6.6 | 29.2 | 55.6 | - | 6.952 | 5.950 |

***PC spines***

| | | | 95% CI | | P-value[†] / Z-score | | |
|---|---|---|---|---|---|---|---|
| **Compartment**[*] | **mean**[**] | **s.e.m** | **Lower** | **Upper** | **exPSD** | **PSD** | **Spine** |
| Spine | 114.0 | 12.9 | 91.0 | 142 | 0.749 | 0.840 | - |
| PSD | 124.0 | 16.5 | 95.6 | 161 | 0.805 | - | -0.564 |
| exPSD | 111.0 | 14.8 | 85.0 | 144 | - | 0.628 | 0.724 |

[*]AZ = active zone, exAZ = extra AZ, PSD = postsynaptic density, exPSD = extra PSD

[**]Marginal means estimated from generalized mixed-effects models

[†]P-values are calculated using mutiple comparison with Tukey method
